# Virtual patient analysis identifies strategies to improve the performance of predictive biomarkers for PD-1 blockade

**DOI:** 10.1101/2024.05.21.595235

**Authors:** Theinmozhi Arulraj, Hanwen Wang, Atul Deshpande, Ravi Varadhan, Leisha A. Emens, Elizabeth M. Jaffee, Elana J. Fertig, Cesar A. Santa-Maria, Aleksander S. Popel

**Affiliations:** Department of Biomedical Engineering, Johns Hopkins University School of Medicine, Baltimore, MD 21205, United States; Department of Oncology, Sidney Kimmel Comprehensive Cancer Center, Johns Hopkins University School of Medicine, Baltimore, MD 21205, United States; Convergence Institute, Johns Hopkins University School of Medicine, Baltimore, Maryland; Bloomberg Kimmel Immunology Institute, Johns Hopkins University School of Medicine, Baltimore, Maryland; Independent consultant; Department of Applied Mathematics and Statistics, Whiting School of Engineering, Johns Hopkins University, Baltimore, Maryland

**Keywords:** Predictive biomarkers, immunotherapy, PD-1 blockade, metastatic triple-negative breast cancer, multivariate biomarkers, early on-treatment biomarkers, non-invasive biomarkers, precision medicine

## Abstract

Patients with metastatic triple-negative breast cancer (TNBC) show variable responses to PD-1 inhibition. Efficient patient selection by predictive biomarkers would be desirable, but is hindered by the limited performance of existing biomarkers. Here, we leveraged in-silico patient cohorts generated using a quantitative systems pharmacology model of metastatic TNBC, informed by transcriptomic and clinical data, to explore potential ways to improve patient selection. We tested 90 biomarker candidates, including various cellular and molecular species, by a cutoff-based biomarker testing algorithm combined with machine learning-based feature selection. Combinations of pre-treatment biomarkers improved the specificity compared to single biomarkers at the cost of reduced sensitivity. On the other hand, early on-treatment biomarkers, such as the relative change in tumor diameter from baseline measured at two weeks after treatment initiation, achieved remarkably higher sensitivity and specificity. Further, blood-based biomarkers had a comparable ability to tumor- or lymph node-based biomarkers in identifying a subset of responders, potentially suggesting a less invasive way for patient selection.

## Introduction

Immune checkpoint inhibitors (ICIs) demonstrate variable efficacy among patients, highlighting the need for personalized treatment approaches. While a fraction of patients achieve long-lasting anti-tumor responses, the vast majority of patients show poor responses to ICIs in many cancer types^1–5^. Due to the incidence of immune-related adverse events^6, 7^, it is critical to ensure that the benefits of ICIs outweigh the risks; thus, there is a need to prospectively identify the patients expected to respond well to ICIs. Consequently, there is a large focus on identifying predictive biomarkers that assist in patient selection for ICI treatment^8^. Predictive biomarkers, such as programmed cell death-ligand 1 (PD-L1) expression, tumor mutational burden (TMB), tumor infiltrating lymphocytes (TILs), neoantigen burden, mismatch repair deficiency/microsatellite instability-high (dMMR/MSI-H), T cell clonality and specific gene mutations^9^, have been recognized. These biomarkers are proven to be applicable to different extent in different cancer types and indications, but in general have limited predictive power. The modest performance of established biomarkers is primarily attributed to the multifaceted impact of the tumor microenvironment (TME) on the therapeutic immune response. The intricate network of cross-talk between distinct cell types and the activity of highly convoluted feedback mechanisms^10, 11^ create a harsh environment for immune effectors to function. Moreover, intra-tumoral, inter-tumoral and inter-individual heterogeneities, at both the spatial and temporal scales, impose additional challenges in the identification of highly effective predictive biomarkers. This complex environment necessitates a systems-level understanding^12–14^.

In recent years, there had been numerous attempts to tackle these challenges and to improve patient selection. One approach aims at combining multiple biomarkers to enhance performance compared to individual biomarkers^15^. TMB and a T-cell inflamed gene expression profile, when jointly assessed, had a superior ability to identify responders to PD-1 inhibition compared to individual biomarkers^16^. Apart from biomarkers assessed before treatment initiation (baseline biomarkers), dynamic changes in biomarker levels at early stages of treatment also has the potential to determine whether a patient is likely to benefit from continued treatment. In a clinical trial with durvalumab and tremelimumab in metastatic breast cancer, a significant expansion in a subset of T cells and increases in CD8, granzyme and perforin expression levels were observed during treatment in responders^5^. Non-invasive liquid-biopsy based biomarkers, such as circulating tumor DNA, circulating tumor cells, immune cell counts or cytokine levels in plasma, are also being explored as an alternative to invasive tissue biopsy-based biomarkers^17^. Advancements in artificial intelligence (AI), machine learning (ML) and bioinformatics together with the increased availability of omics data have enabled the identification of multi-omics-based biomarkers to predict treatment responses^18–20^. Despite these parallel efforts, there is no consensus on the best strategy to improve the performance of predictive biomarkers.

Triple-negative breast cancer (TNBC), which is an aggressive subtype of breast cancer characterized by the absence of estrogen and progesterone receptors and the lack of HER2 overexpression, has emerged as an attractive target for ICIs^21–23^. However, single-agent ICIs have very low response rates in metastatic TNBC^4, 24–27^, leading to the exploration of combination therapies^28–30^. The combination of pembrolizumab and chemotherapy is approved for patients with PD-L1-positive metastatic TNBC in the first-line setting^28^. However, a significant fraction of patients with TNBC have PD-L1 negative tumors, and treatment options are limited for patients who experience disease progression after one or more lines of systemic treatment in the metastatic setting. Thus, there is an unmet clinical need to identify patients likely to respond to single-agent ICIs and to determine the best immunotherapy combinations or alternative therapies for non-responders^23^. Exploratory analysis of patients treated with pembrolizumab (anti-PD-1 antibody) monotherapy in KEYNOTE-086 trial^24, 25^ suggested an association of objective response rates with biomarkers such as PD-L1 combined positive score, CD8, stromal TILs, TMB and T-cell inflamed gene expression profile^31^. KEYNOTE-119 data^26^ suggested that while PD-L1 expression on immune cells can independently predict response, addition of PD-L1 expression on tumor cells could improve the predictive value^32^. While an in-depth analysis of biomarker candidates including the cellular and molecular components of the tumor microenvironment would help inform future clinical studies for metastatic TNBC, the limited availability of patient-level data associated with adequate biospecimens for complex testing remains a challenge for exploratory biomarker analyses.

Partially synthetic data generated using bespoke mathematical models informed with real data is an attractive alternative for hypothesis generation and decision making in health care^33^. A number of mathematical models have been developed for exploratory analyses to gain novel insights into treatment outcomes^34–36^ and for investigating the implications of varying trial designs^37^. These models are powerful tools for simulating personalized therapies^38–40^. Multi-scale mechanistic models, such as quantitative systems pharmacology (QSP) models that are built and calibrated using a combination of in-vitro, in-vivo, clinical, population level data, and multi-omics data, have enabled the generation of virtual patients to simulate in-silico clinical trials^41–49^. Due to the mechanistic nature and the ability to predict time profiles of cellular and molecular species of interest, QSP model-generated virtual patients provide a wealth of longitudinal patient-level data that can be leveraged for numerous applications, including biomarker testing. Here, we use patient-level data generated using a QSP model of metastatic TNBC^42^ both to identify predictive biomarkers of response to single agent ICI pembrolizumab, and to explore various strategies to enhance the predictive power of biomarker candidates.

## Materials and Methods

### Brief description of QSP model for metastatic TNBC

Our bespoke QSP model for metastatic TNBC^42^ was used to generate partially synthetic data for biomarker testing. Briefly, the QSP model is a multi-scale, compartmental, whole-patient model which accounts for mechanisms associated with the tumor progression, anti-tumor immune responses and pharmacodynamics/pharmacokinetics of ICIs. The QSP model incorporates multiple compartments including primary tumor, metastatic tumors, tumor draining lymph node, central, and peripheral compartments. As lung metastasis is very common in TNBC, two lung metastatic tumor compartments were incorporated. In addition, a “other” metastatic tumor compartment is considered to represent metastases in organs other than the lung. The model considers different steps involved in the cancer immunity cycle^50^, such as the proliferation and death of cancer cells in the tumor compartments resulting in the release of self-antigens and neo-antigens, antigen uptake by antigen-presenting cells and their maturation, activation of helper, cytotoxic and regulatory T cells in the lymph node compartments, trafficking of T cells to the tumor compartment, anti-tumoral activity of effector T cells, pro-tumoral activity of regulatory T cells and exhaustion of effector T cells. The tumor compartments also include additional anti-tumorigenic as well as pro-tumorigenic cell types, such as M1 macrophages, M2 macrophages, myeloid derived suppressor cells and molecular species, such as PD-L1, IL-10, IL-12, CCL2, NO, Arg-I, TGFβ and IFNγ. The activity of cytokine IL-2 in promoting the activation of T cells is considered in the lymph node compartments. The model is built based on the model of a single primary tumor^47^, which comprises 152 ordinary differential equations (ODEs), and 280 parameters. In the current model, the metastatic compartments have the same structure as the primary tumor compartment and most of the parameters, such as the binding rate constants, are shared between the compartments and they have been selected in our previous publications^47, 51^. Parameters such as the cancer cell proliferation rate constant, initial APC density, seeding time of metastatic tumors, recruitment rates of MDSCs and macrophages, polarization rate constant of M1 to M2 macrophages, are compartment specific^42^. In addition to multiple tumor compartments, multiple cancer clones, neo-antigens and T cell specificities are considered that leads to the corresponding increase in the number of variables: 641 ODEs and 737 parameters. A combination of data from in-vitro, in-vivo and clinical studies were used for estimating the parameters of the model, including immuno-genomic data. The model is implemented using the SimBiology toolbox of MATLAB (MathWorks, Natick, MA). Detailed descriptions of the QSP model and parameterization are provided in^42^.

### Virtual patient generation and treatment simulations to generate partially synthetic data

Selected parameters of the QSP model were sampled using Latin hypercube sampling (LHS) to generate virtual patients (i.e., each unique parameter set corresponds to a virtual patient). Physiologic distributions for each of the sampled parameters were estimated in our previous study^42^ using the population-level clinical response data from KEYNOTE-119 trial^26^. Thus, the generated virtual patient cohort is representative of patients with TNBC undergoing pembrolizumab monotherapy as a second or later line therapy in the metastatic setting. To generate initial conditions for treatment simulations (i.e. pre-treatment or baseline cellular and molecular compositions), virtual patients were simulated without the administration of pembrolizumab until at least one of the metastatic tumors attained the target tumor diameter. QSP model variables at the time point where the virtual patients attained the target tumor diameter were determined and were used as initial conditions for treatment simulations. Virtual patients that did not attain the target tumor diameter were considered unphysiological and were excluded from any subsequent analysis. Number of cancer cells in the primary tumor compartment was set to 0, as most patients with metastatic TNBC would have undergone surgical removal of the primary breast tumor at early stages. Seeding time of metastatic tumor compartments were randomly sampled resulting in virtual patients with varying number of tumors (1-3 metastatic tumors per patient). The virtual patient cohort used in this study includes a total of 1635 patients. Virtual patients were classified based on their response statuses as complete/partial response (CR/PR), stable disease (SD) or progressive disease (PD) based on the change in the sum of tumor diameter as per RECISTv1.1^52^. Thus, the partially synthetic data include model-generated time profiles of cellular and molecular species during pembrolizumab treatment of virtual patients with metastatic TNBC.

### Selection of biomarker candidates

Various molecular and cellular species from different compartments of the QSP model were selected as biomarker candidates. This includes a total of 90 biomarker candidates including densities of cell types, concentrations of cytokines or soluble mediators, ratios of different cellular populations, expression of immune checkpoint molecules, clonality measures such as the richness, evenness, and diversity^53, 54^ of T cells or cancer cells and the tumor diameter. The complete list of biomarker candidates selected is provided in **Table S1**. Values of the selected biomarker candidates were extracted from all virtual patients at baseline (pre-treatment). In subsequent analysis, responders include patients with SD for at least 24 weeks and patients with CR/PR, whereas non-responders refer to patients with PD.

### Cutoff-based biomarker testing algorithm

A cutoff-based biomarker testing algorithm was used to evaluate and rank biomarkers. This method is based on the algorithm described in^42^, and has been modified to improve the computational speed, robustness and predictability in unseen dataset. An outline and a schematic illustration of the optimized algorithm is provided in **Fig S1**. In-silico virtual patients generated were randomly split into training and test sets with 1145 (70%) and 490 patients (30%), respectively. Training set was used to identify optimal cutoff values for each biomarker candidate and the performance of the identified optimal cutoff was then validated using the test dataset. To identify the optimal cutoff, the following steps are followed for each biomarker candidate: First, 8 quantiles of the population distribution of the biomarker candidate were selected as potential cutoff values. Virtual patient subsets were generated by identifying patients with biomarker candidate levels greater than or equal to each selected cutoff resulting in 8 non-mutually exclusive virtual patient subsets. To account for biomarker candidates having negative correlation with responses (i.e., lower level of a biomarker candidate associated with better response), we also generated virtual patient subsets with biomarker candidate levels less than the selected cutoffs. This results in a total of 16 virtual patient subsets (8 cutoffs x 2 conditions (greater than or equal to the cutoff; less than the cutoff)). Subsets with less than 20 virtual patients (∼ 2% of total patients in the training set) were discarded to avoid artifacts due to low number of patients. The virtual patient subsets were then evaluated with performance measures **(see section Performance measures)** to identify the optimal cutoff and condition for each biomarker candidate. Biomarker candidates are ranked based on the performance measure calculated for the optimal cutoffs and conditions.

### Performance measures

To evaluate the performance of biomarker candidates, two different metrics are used. The first measure, response probability is defined as the fraction of responders among the virtual patients in the selected subset. The second measure, RIS or responder inclusion score is a metric that we previously introduced^42^ to increase the selection of responders from the entire patient cohort (i.e., to increase sensitivity as well as specificity). RIS is calculated as the fraction of responders selected from the entire cohort minus the fraction of non-responders selected from the entire patient cohort.

### Feature selection and multivariate biomarkers

For identifying biomarker combinations, an initial feature selection step was applied to select important features, followed by testing combinations of selected candidates with the cutoff-based biomarker testing algorithm. This step was included to reduce the computational cost in testing multiple cutoffs for biomarker combinations. Training dataset with biomarker candidate levels and class labels (responders or non-responders) of 1145 virtual patients was used as the input for feature selection. Feature selection was performed with random forest classifier using the recursive feature selection algorithm of the sklearn toolkit in python. F1 score, which is the harmonic mean of precision and recall, was used as the metric to identify the optimal number of features and select important features or biomarker candidates. Then, cutoff-based biomarker testing algorithm was applied on the selected biomarker candidates to test panels of double, triple, and quadruple biomarkers **(Fig S1)**. Multiple panels combining selected biomarker candidates were generated, and the cutoff-based biomarker algorithm was used to test combinations of cutoffs for biomarkers in each panel. While all combinations of cutoffs for biomarkers in each panel are tested, the conditions (greater than or equal to the cutoff; less than the cutoff) are chosen based on the optimal condition from the single biomarker analysis. Following the steps of single biomarker testing, virtual patient subsets were generated that satisy the cutoff criteria of biomarkers in each panel. Then, the optimal set of cutoffs for each biomarker panel was identified by evaluating performance measures. Performance of single biomarkers were then compared with multivariate biomarker panels.

### On-treatment biomarkers

To compare the predictive power of pre-treatment biomarker candidates with early on-treatment biomarkers, we extracted the biomarker candidate levels at day 15 and day 30 after treatment initiation. Relative change of biomarker candidate levels on day 15 or day 30 with respect to baseline were also calculated. As described for pre-treatment biomarker candidates, cutoff-based biomarker testing algorithm was applied without feature selection for single biomarkers and with random forest-based feature selection for multivariate biomarker panel evaluation. Performance of pre-treatment biomarkers were compared with relative change after treatment initiation (“rel. change day 15” and “rel. change day 30”) as well as single time point on-treatment biomarkers (“day 15” and “day 30”).

### Biomarkers from different compartments

Biomarker candidates were classified as lymph node-, tumor- or blood-based biomarkers based on their compartment of origin in the QSP model. Blood-based biomarkers refer to biomarker candidates from the central compartment of virtual patients. For lymph node- and metastatic tumor-based biomarker candidates, corresponding quantities from the lymph nodes and metastatic tumors of each patient were averaged. Apart from the cellular and molecular characteristics of tumor, tumor diameter that is routinely determined by computed tomography imaging in clinical practice was also considered as a tumor-based biomarker. Collectively, biomarker candidates selected included 14, 16 and 60 candidates associated with blood, lymph node and tumor, respectively.

### In-silico biomarker validation trials

To conduct in-silico biomarker validation trials, a second virtual patient cohort including a total of 829 patients was generated as discussed in the section **Virtual patient generation and treatment simulations to generate partially synthetic data**. However, a different seed was used for random number generation and thus, the generated virtual patient cohort is distinct from the first cohort with 1635 patients that was used for the identification of biomarker candidates. Representative biomarkers or panels of non-invasive biomarkers with high response probability and RIS were selected. Virtual patients in the unseen second cohort were stratified based on selected biomarkers and in-silico virtual trials were conducted to predict the trial outcomes in biomarker selected population.

### Software and data availability

QSP model virtual patient generation and in-silico clinical trials were performed using the MATLAB code that can be accessed via Mendeley data (https://data.mendeley.com/datasets/r46rk4vwdv/1). R software was used for cutoff-based biomarker testing and visualization of the results. random forest-based feature selection was performed using sklearn toolkit of python.

## Results

### Pre-treatment individual biomarkers have modest performance

The workflow of this study involved generating virtual patients from a QSP model, biomarker testing using partially synthetic data derived from virtual patients and evaluating various strategies for improving the predictive power of biomarker candidates as depicted in **Fig 1a**. We used the QSP model for metastatic TNBC^42^ to generate 1635 virtual patients by Latin hypercube sampling of selected parameters. QSP model building, and virtual patient generation utilized population-level clinical data from KEYNOTE-119 trial^26^, and immune cell composition estimates from RNA-seq data^55^ and mechanistic details gathered from several in-vivo/in-vitro data. We conducted in-silico clinical trial of pembrolizumab monotherapy, resulting in the generation of partially synthetic or in-silico, patient-level, time course data. In-silico trial outcomes adequately recapitulated population characteristics of real patient cohort from KEYNOTE-119 clinical trial^26, 56^ including response rate, duration of response, time to response and proportion of patients with lung metastases **(Fig 1b).**

**Figure 1:**
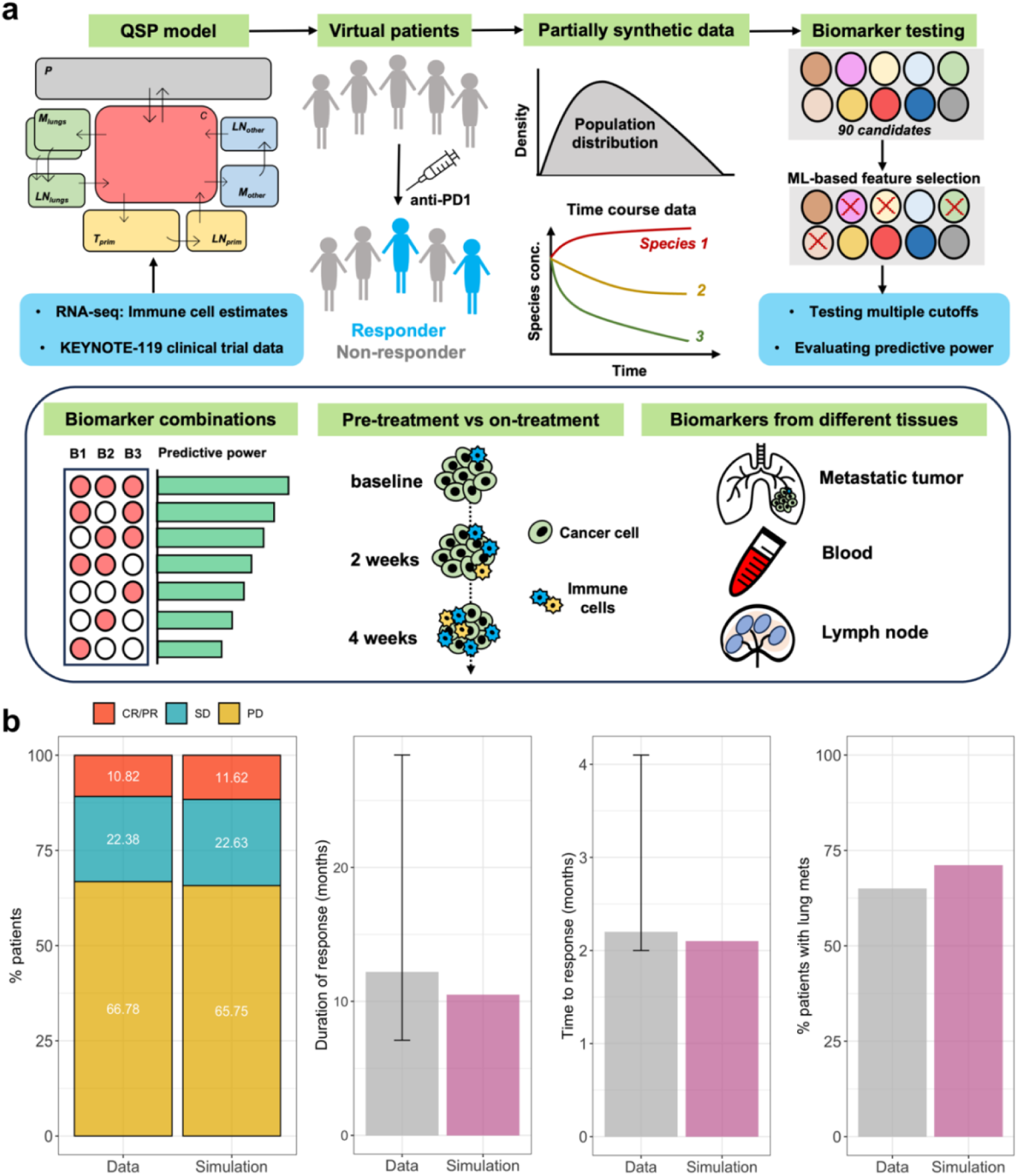
**a)** Workflow of the study. QSP model informed with RNA-seq, and clinical data was used to generate virtual patients with metastatic TNBC. In-silico clinical trial of pembrolizumab was conducted resulting in the generation of partially synthetic data such as population distribution and longitudinal data of various species. For biomarker testing, 90 candidates including various model species were selected. ML-based feature selection was used to pre-filter biomarker candidates, followed by evaluating the performance by testing various cutoffs for selected biomarkers. Various strategies to improve the performance compared to single pre-treatment biomarkers were tested such as by combining multiple biomarkers and assessing on-treatment biomarkers. Finally, the performance of biomarkers from different tissues (blood, lymph node and metastatic tumor) were compared. **b)** Comparison of in-silico trial outcomes with clinical data from KEYNOTE-119 trial for response rates, duration of response, time to response and percentage of patients with lung metastases. KEYNOTE-119 data^26, 56^ was renormalized by excluding non-evaluable patients for comparison with simulations.

Generated virtual patients were randomly split into training and test sets with 1145 patients in training and 490 patients in the test set. We extracted the pre-treatment or baseline levels of 90 biomarker candidates **(Table S1)** from virtual patients in the training set and leveraged the modified version of cutoff-based biomarker testing algorithm to identify the cutoffs that resulted in the optimal selection of responders using our first metric response probability. Response probability, which is defined as the fraction of responders among the subset of patients selected, has a theoretical maximum value of 1, when the subset exclusively contains only responders. Thus, specificity, which is defined as the proportion of non-responders correctly excluded from selection, is higher when the response probability is high. On the other hand, a high response probability does not necessarily imply a high sensitivity, which is defined as the proportion of positive patients correctly selected by biomarkers. When the biomarker candidates were sorted based on the response probability **(Fig 2a)**, antigen presenting cells (APCs) in lymph nodes, which attained a response probability of 0.74, was the top ranked biomarker. This was followed by the level of regulatory T cells (Tregs) and the fraction of cytotoxic T cells (Tcyt) among total T cells in lymph nodes. Top 15 biomarkers based on response probability also included the tumor diameter, clonality measures of T cells and cancer cells, and PD-L1 expression in the tumor compartment. Virtual patient subsets selected by the best single biomarker based on response probability included 5% of non-responders and only 30% of responders from the whole patient cohort **(Fig 2a)**, thus showing modest specificity but low sensitivity.

**Figure 2:**
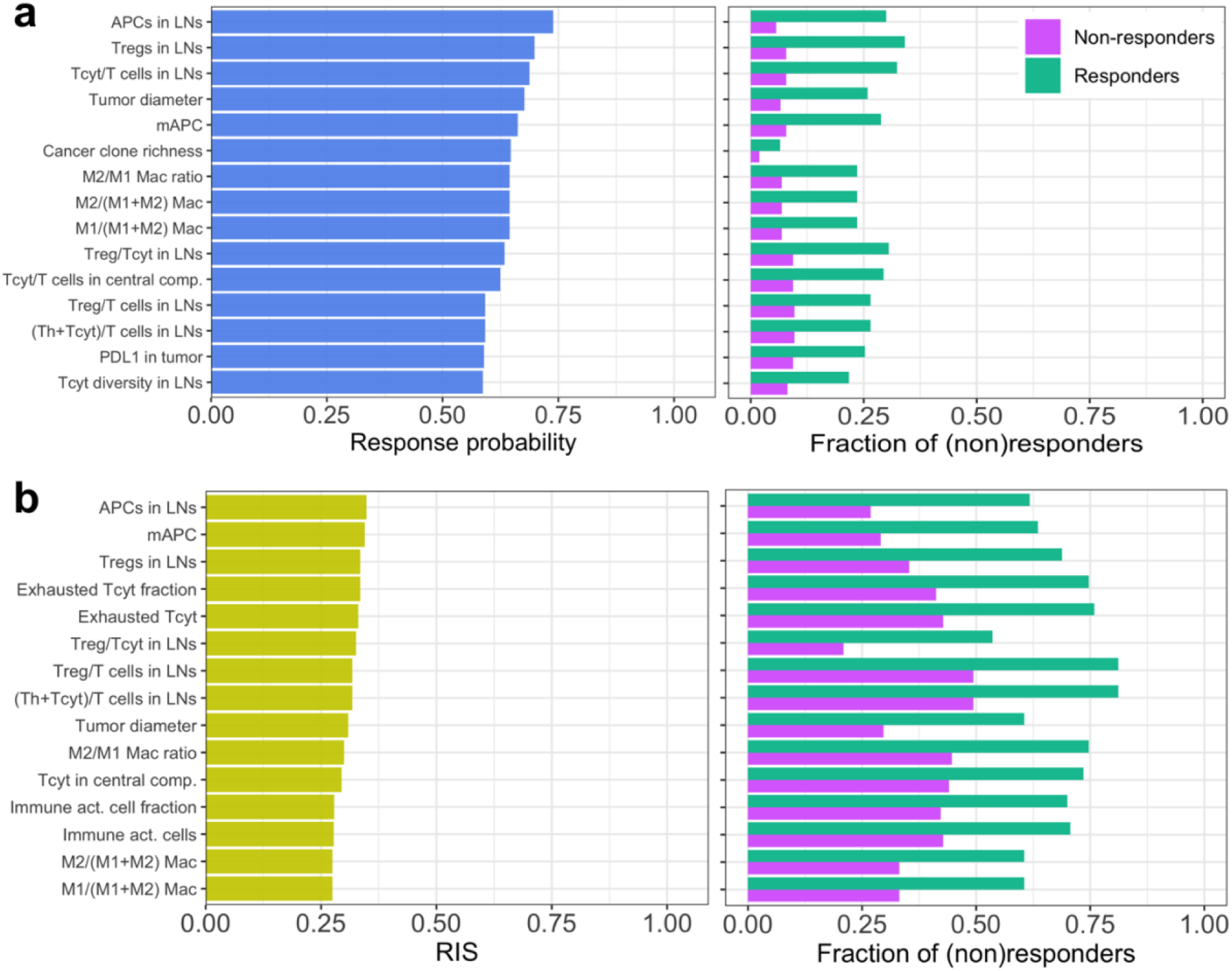
Pre-treatment single biomarker analysis. Top 15 pre-treatment predictive biomarkers identified based on metrics **a)** response probability and **b)** responder inclusion score or RIS. Panels on the right show the fraction of responders and non-responders from the whole patient cohort selected by corresponding biomarkers.

Conversely, we also tested the negative predictive value of pre-treatment single biomarker candidates defined as the ability of biomarker candidates to identify non-responders rather than identifying responders. 1-response probability was used as the performance measure in this case. Like positive predictive biomarkers, single pre-treatment biomarkers had only modest ability to identify non-responders as the top biomarker candidates were only able to identify less than 25% of non-responders from the entire cohort and in addition, ended up selecting a small fraction of responders **(Fig S2).**

The second metric RIS or responder inclusion score, has a theoretical maximum value of 1, when the sensitivity and specificity are both 100%. When the biomarker testing algorithm was implemented considering RIS as the metric, the maximum RIS attained was 0.35 for APCs in lymph nodes **(Fig 2b)**. The virtual patient subset selected based on this biomarker had 62% of responders from entire cohort implying high sensitivity compared to top biomarkers based on response probability. However, a larger percentage of non-responders (27%) were also included implying a low specificity. Interestingly, 10 biomarkers were shared between the top 15 biomarkers based on RIS and response probability.

### Cutoffs determine trade-off between sensitivity and specificity of predictive biomarkers

We examined the changes in the performance of biomarkers for varying cutoffs **(Fig 3)**. For most biomarker candidates, response probability and RIS varied considerably for different cutoffs. Despite the presence of shared biomarkers between top 15 biomarkers based on RIS and response probability **(Fig 2)**, the cutoff value that resulted in maximum RIS or response probability were different for several biomarkers **(Fig 3)**. For instance, for tumor diameter, maximum response probability was attained at a cutoff value of 2.2 cm, but maximum RIS was attained at a cutoff value of 1.7 cm **(Fig 3a)**. Further, with varying cutoffs, the performance of biomarkers varied non-monotonically, especially for RIS **(Fig 3)**. These results suggest that the cutoff values are an important determinant of the tradeoff between RIS and response probability.

**Figure 3:**
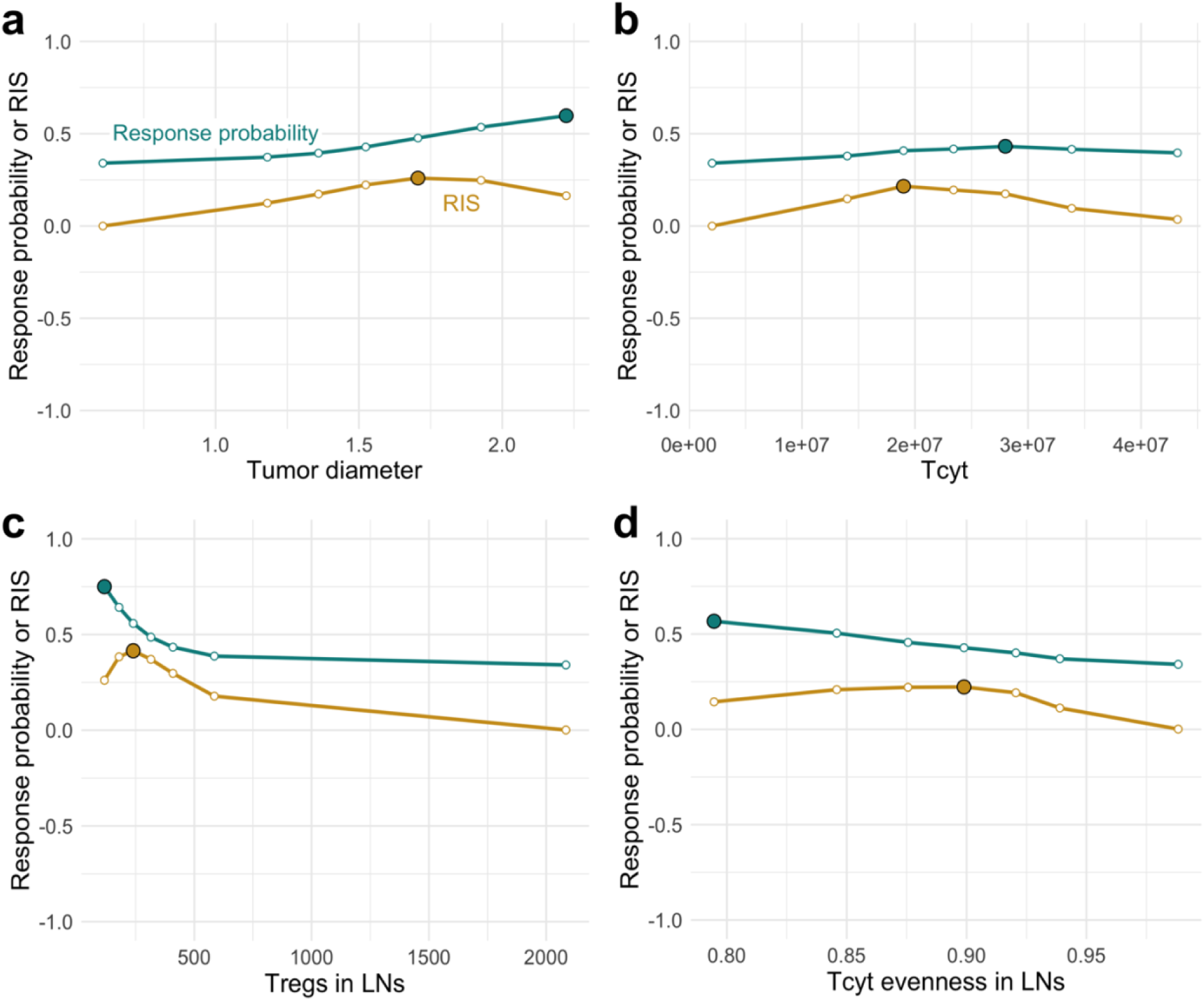
Response probability or responder inclusion score (RIS) for varying cutoffs of **a)** Tumor diameter in centimeters, **b)** Tcyt density in the tumor (cells/ml), **c)** Treg density in lymph nodes (cells/mm^3^) and **d)** Tcyt evenness in lymph nodes. Highlighted point (solid circle with black border) represents the biomarker cutoff that resulted in maximum response probability or RIS.

### Combining biomarkers improves the specificity of pre-treatment biomarkers

We performed feature selection by recursive feature elimination with random forest classifier and generated multivariate biomarker panels, such as double, triple, and quadruple combinations of selected biomarker candidates. Random forest selected biomarker candidates are listed in **Table S1**. Combinations of biomarkers showed an increase in response probability **(Fig 4a)**. The extent of increase in response probability varied for different biomarker combinations, and multivariate biomarkers with a response probability of 1 or 100% specificity were also seen **(Fig 4a)**. RIS increased for multivariate biomarkers compared to single biomarkers, but the increase was modest **(Fig 4b)**. The distribution of response probability and RIS showed a prominent increase in the performance for panels with two biomarkers compared to single biomarkers, but only minimal changes were seen for panels with three or four biomarkers compared to panels with two biomarkers **(Fig 4c and 4d)**.

**Figure 4:**
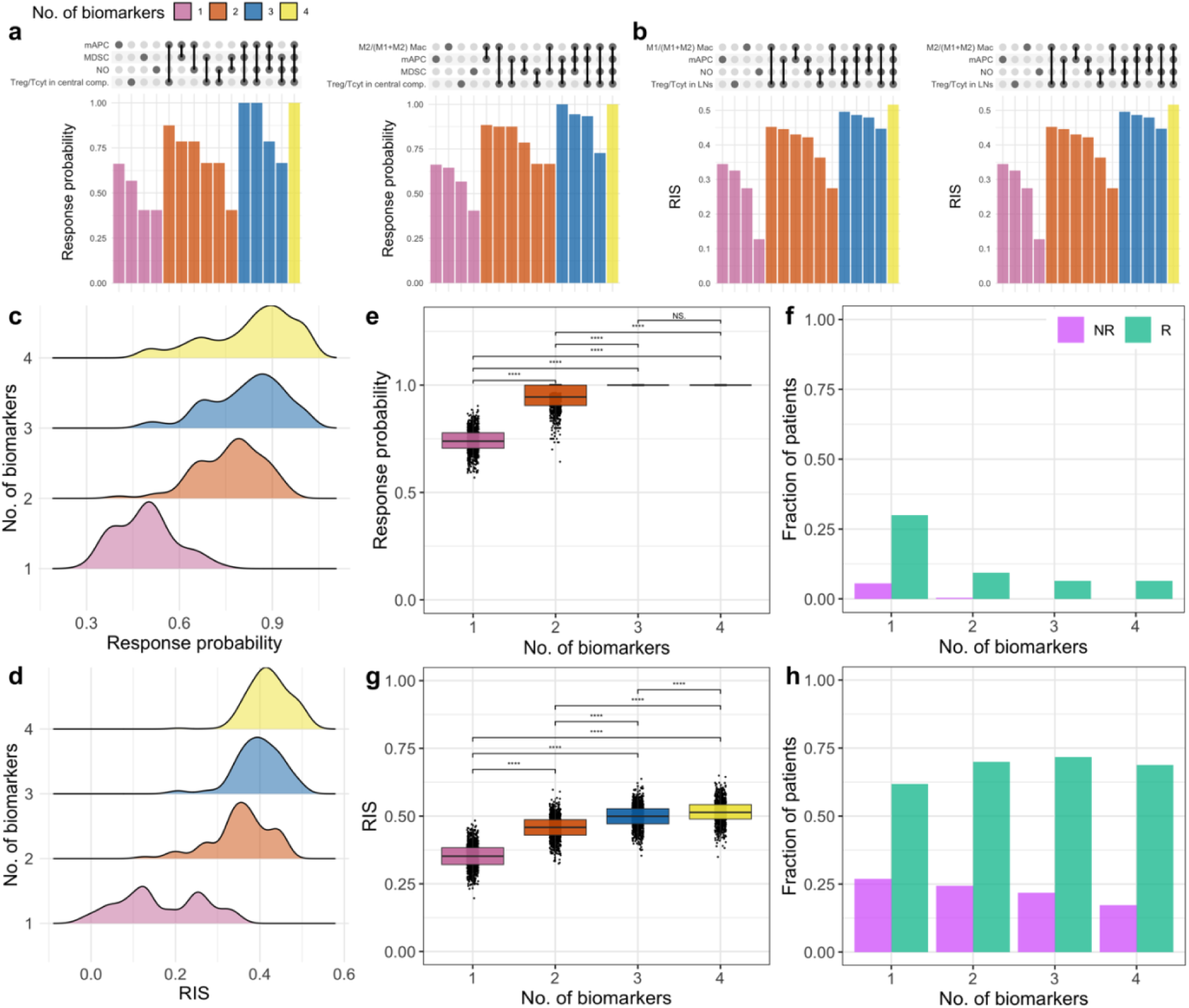
Multivariate pre-treatment biomarkers. **a** and **b)** UpSet plots of response probabilities **(a)** and RIS **(b)** for representative single and multivariate biomarkers. Bars represent the response probabilities/RIS of biomarker combinations indicated at the top of corresponding bars as highlighted circles connected by lines. **c** and **d)** density plots of response probability and RIS for tested single and multivariate biomarkers. **e** and **g)** Response probabilities **(e)** and RIS **(g)** of the best multivariate biomarkers compared with the best single biomarker. **f** and **h)** Fraction of responders and non-responders among entire patient cohort selected by best multivariate and single biomarkers based on response probability **(f)** or RIS **(h)**. Statistical significance was calculated using the Wilcoxon test (NS: p > 0.05; ****: p<=0.0001).

The best biomarker panels with two, three and four biomarkers among the tested panels attained response probabilities close to 1 **(Fig 4e)**. While a notable fraction of patients selected by the top single biomarker had progressive disease, selection of non-responders (or false positives) was reduced or eliminated in the best multivariate biomarker panels **(Fig S3a)**. For the best triple and quadruple biomarkers, a large fraction of patients selected had complete or partial responses **(Fig S3a)**. This implies that when patients are selected by multivariate biomarkers, a very high response probability could be achieved. However, the fraction of responders from the entire patient cohort selected was low even with multivariate biomarkers **(Fig 4f)**. Thus, a remarkable increase in specificity with pre-treatment biomarker combinations comes at the cost of reduced sensitivity (i.e., increased false negatives). When RIS of best single biomarker was compared with biomarker combinations, there was an increase from 0.35 to 0.52 **(Fig 4g)**. When compared to multivariate biomarkers based on response probability, biomarkers identified based on RIS were able to correctly identify a much larger proportion of responders from the entire cohort (higher sensitivity) at the cost of reduced specificity **(Fig 4h; Fig S3b)**. Thus, while combinations of pre-treatment biomarkers were able to increase specificity, there was a limited ability to simultaneously achieve both high specificity as well as sensitivity.

### On-treatment biomarkers improve sensitivity as well as specificity compared to pre-treatment biomarkers

To test whether on-treatment biomarkers can overcome the limitations of pre-treatment biomarkers and improve sensitivity as well as specificity, we extracted biomarker candidate levels from virtual patients at day 15 and day 30 after the initiation of pembrolizumab treatment. We also calculated relative change in biomarker levels on day 15/30 with respect to pre-treatment values. UMAP dimensionality reduction with pre-treatment biomarker candidate levels showed no clear separation of patients with different response statuses **(Fig 5; baseline)**. For on-treatment biomarkers, patients with complete/partial responses were confined and appeared to form a cluster **(Fig 5),** suggesting that on-treatment biomarkers likely contain more information, that allows a better separation of patients with different response statuses.

**Figure 5:**
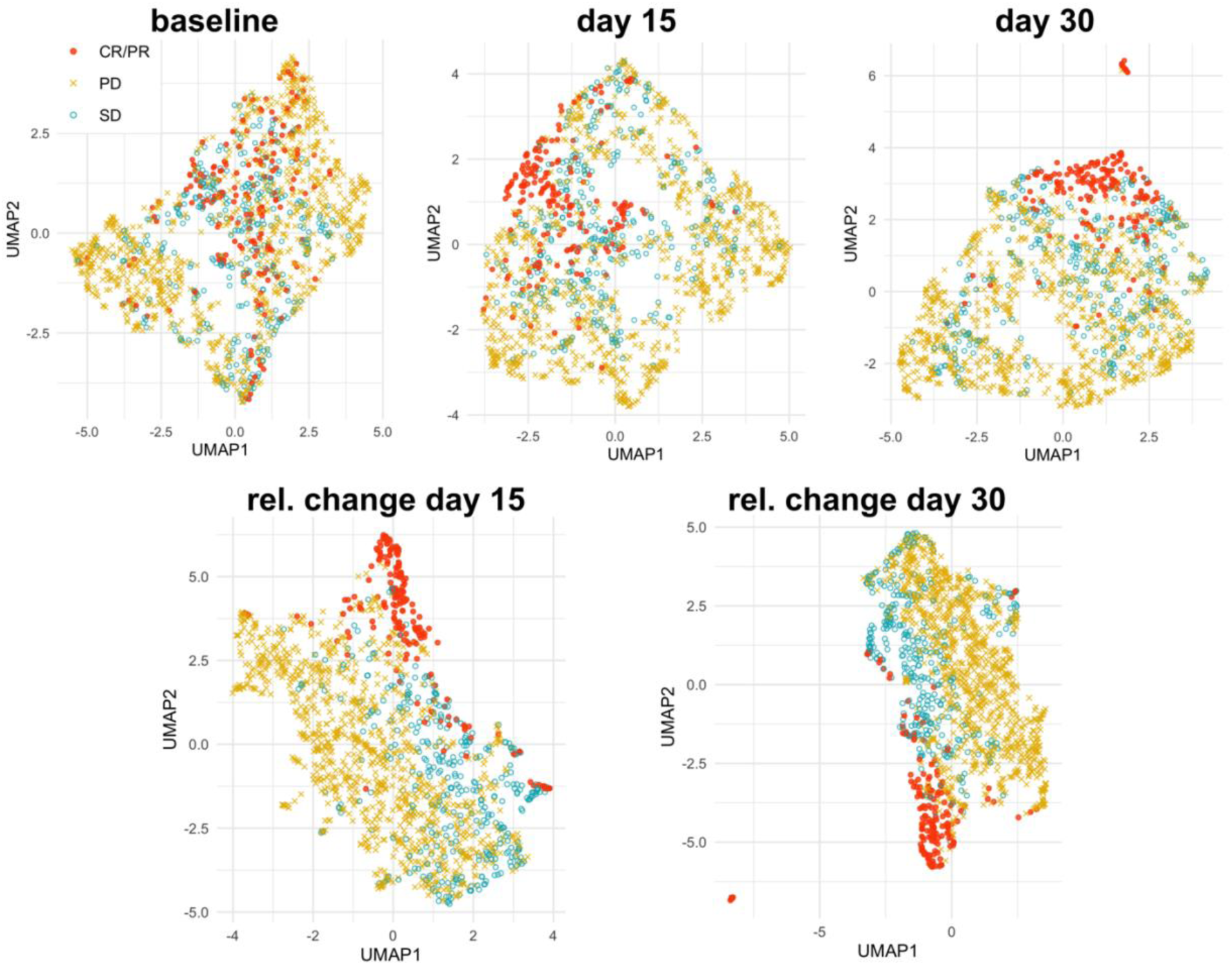
Dimensionality reduction with Uniform Manifold Approximation and Projection for biomarker candidate levels at baseline, day 15 or day 30 after treatment initiation and relative change on day 15 or day 30 with respect to baseline. Colors and shapes indicate response statuses based on RECISTv1.1. CR/PR: Complete/Partial response; SD: Stable disease; PD: Progressive disease.

We then compared the predictive power of single biomarker candidates at baseline, day 15, day 30 and relative change at day 15 or day 30 with respect to baseline. For relative change on day 15 with respect to baseline, more than 50% of biomarker candidates showed an increase in response probability compared to performance of the same biomarker candidate at baseline **(Fig 6a)**. Thus, several single biomarker candidates, such as helper T cell density in the central compartment, arginase I, CCL2 and macrophage density in the tumor, tend to show an increase in specificity when the relative change with respect to baseline level is examined. Biomarker candidates, such as the overall PD-L1 expression in the tumor, fraction of M2 macrophages among total macrophages in the tumor, showed a decrease in response probability when the relative change with respect to baseline is considered compared to baseline **(Fig 6a)**. Similarly, biomarker candidates, such as fraction of TILs the in tumor, fraction of M2 macrophages among total macrophages in the tumor, showed a decrease in RIS when the dynamic change with respect to baseline was considered. Interestingly, some biomarkers, such as Tcyt density in lymph nodes and Treg fraction in tumor, showed a remarkable increase in RIS at 2 weeks after treatment initiation **(Fig 6b)**. Like relative change on day 15, biomarker candidates showing either an increase or decrease in performance were observed when tested at day 15, day 30 or relative change at day 30 with respect to baseline **(Fig S4)**. These results show that while many biomarker candidates show an increased performance when measured after treatment initiation, there is also a notable fraction of biomarker candidates that show a decrease in performance. Therefore, the PD-1 inhibition could increase or decrease the overlap in biomarker distributions of responders and non-responders.

**Figure 6:**
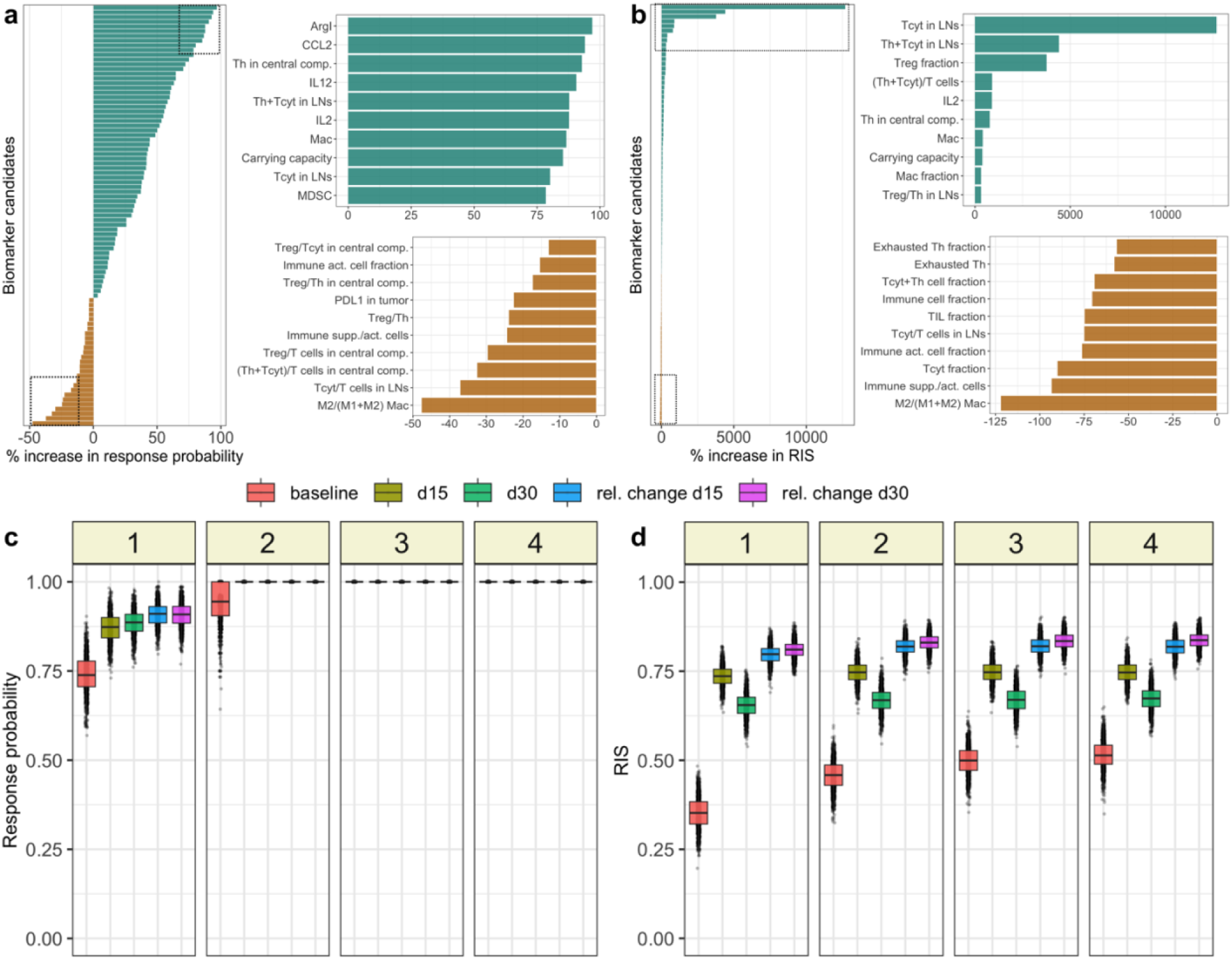
**a** and **b)** Percentage change in response probability **(a)** or RIS **(b)** of relative change in biomarker levels at day 15 from baseline, compared to the performance at baseline. Each horizontal bar corresponds to a single biomarker candidate. Panels on the right show the top 10 biomarker candidates with most increase (green) or decrease (golden yellow) in response probabilities/RIS. **c** and **d)** Comparison of response probabilities **(c)** and RIS **(d)** of best single or multivariate biomarkers for pre-treatment and on-treatment biomarkers. Colors represent different time points of biomarker measurements. Numbers on the top indicate the number of biomarkers considered in each panel. Black dots represent the results of bootstrapping.

We generated multivariate biomarker panels for on-treatment biomarkers by combining feature selection with cutoff-based biomarker testing and compared the best single and multivariate biomarkers identified. Among the best single biomarkers, M2/M1 macrophage ratio which was the best on-treatment biomarker at day 15 had a higher response probability 0.87 compared to 0.74 for the best baseline biomarker **(Fig 6c)**. In addition, relative change in biomarker values on day 15 or day 30 with respect to baseline had a higher response probability compared to biomarker levels at single timepoints after treatment initiation (i.e., on day 15 or day 30). While many single on-treatment biomarkers were superior to baseline biomarkers **(Fig S5 and S6)**, only minimal changes in the distribution of response probability and RIS were observed from two to four on-treatment biomarker combinations **(Fig S7)** as observed with pre-treatment biomarkers. However, the best multivariate baseline biomarkers were able to achieve a response probability of 1 or 100% specificity like multivariate on-treatment biomarkers **(Fig 6c; Fig S8)**. Unlike response probability, best single and multivariate on-treatment biomarkers consistently had higher RIS compared to baseline biomarkers **(Fig 6d; Fig S9)**. These results suggest that tested baseline biomarkers even when combined have lower performance than the best on-treatment biomarkers in simultaneously achieving a good balance between sensitivity as well as specificity, despite the ability to achieve high specificity.

### Non-invasive biomarkers predictive of response

We sought to compare the performance of blood-based biomarkers with tissue-based biomarkers. To this end, we classified biomarker candidates based on the compartment of origin in the QSP model. We tested multivariate biomarker panels of candidates from specific compartments using feature selection followed by cutoff-based biomarker testing. Among single pre-treatment biomarkers, APC density and Treg density from lymph nodes had response probabilities 0.74 and 0.69, respectively, that were higher compared to biomarkers from blood or tumor **(Fig 7a)**. Interesting, tumor diameter, which is a tumor-based biomarker that can be non-invasively (radiologically) measured, had a response probability 0.67. Irrespective of the compartment or timepoint, biomarker panels with three or four biomarkers were able to achieve the highest possible response probability **(Fig 7a and 7c; Fig S10)**. Thus, combinations of pre-treatment or on-treatment blood-based biomarkers offer a non-invasive way to achieve high specificity.

**Figure 7:**
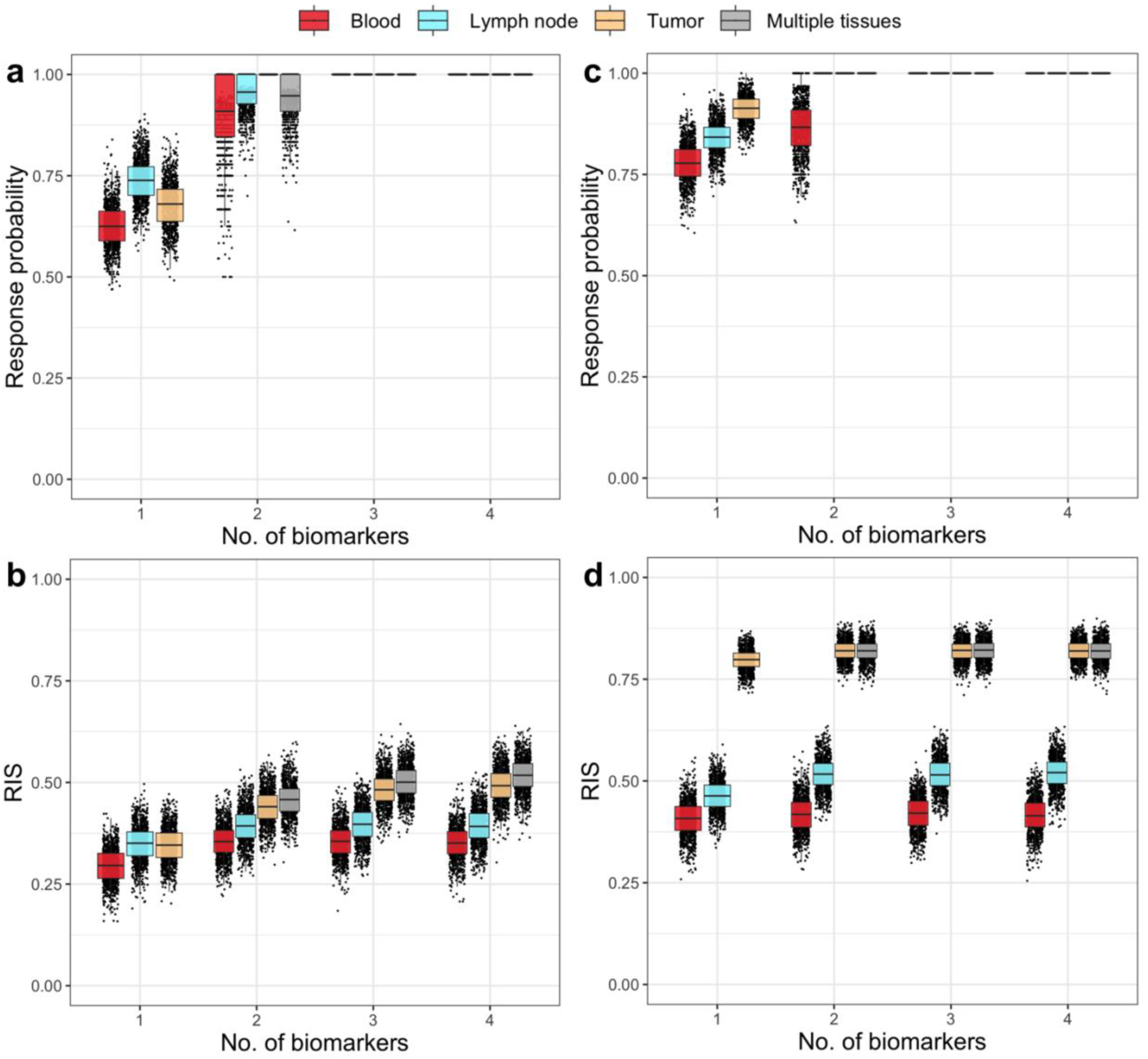
**a** and **b)** Response probabilities and RIS for baseline or pre-treatment biomarkers from different compartments (blood, lymph node or metastatic tumor). **c** and **d)** Response probabilities and RIS of relative change in biomarker levels at day 15 with respect to baseline. Colors indicate biomarkers/biomarker combinations from each compartment. “Multiple tissues” represent combinations of biomarkers from multiple compartments. Black dots represent the results of bootstrapping.

When the ability of biomarkers in simultaneously achieving high sensitivity and specificity (high RIS) was compared between different compartments, best tumor-based biomarkers performed better than the tested blood-based, or lymph node-based biomarkers **(Fig 7b and d; Fig S10)**.

### Biomarker validation with in-silico trials of independent cohort

Having identified biomarker candidates for achieving high specificity alone or a good balance between specificity and sensitivity, we generated an independent patient cohort and tested our predictions by conducting in-silico trials of pembrolizumab with selected patient populations. In the unseen in-silico patient cohort, the model-predicted objective response rate and disease control rate were 10.74% and 32.81%, respectively (in biomarker identification cohort: ORR=11.62%, DCR=34.25%). This simulation corresponds to the unselected patient cohort as in the KEYNOTE-119 trial^26^. As a representative of non-invasive biomarker panel with high specificity, we selected the pre-treatment biomarker combination, density of Tregs and diversity of Tcyt cells, both from the central compartment which represents the blood. Among 829 virtual patients generated, only 18 (2.2%) patients satisfied the biomarker criteria (density of Tregs in central comp. < 345.508 and Tcyt diversity in central comp. < 1.712) and were selected for the in-silico trial. Distribution of selected biomarker levels in the virtual patient cohort is provided in **Figure S11**. In-silico trial in the selected patient cohort showed an objective response rate (ORR) of 22.23%, a disease control rate (DCR) of 77.79%, and a median duration of response 10.5 months **(Fig 8a)**. All virtual patients with CR/PR had a response duration >= 6 months. Furthermore, the in-silico trial showed a shrinkage of multiple metastases in a fraction of virtual patients with multiple tumors **(Fig 8a)**.

**Figure 8:**
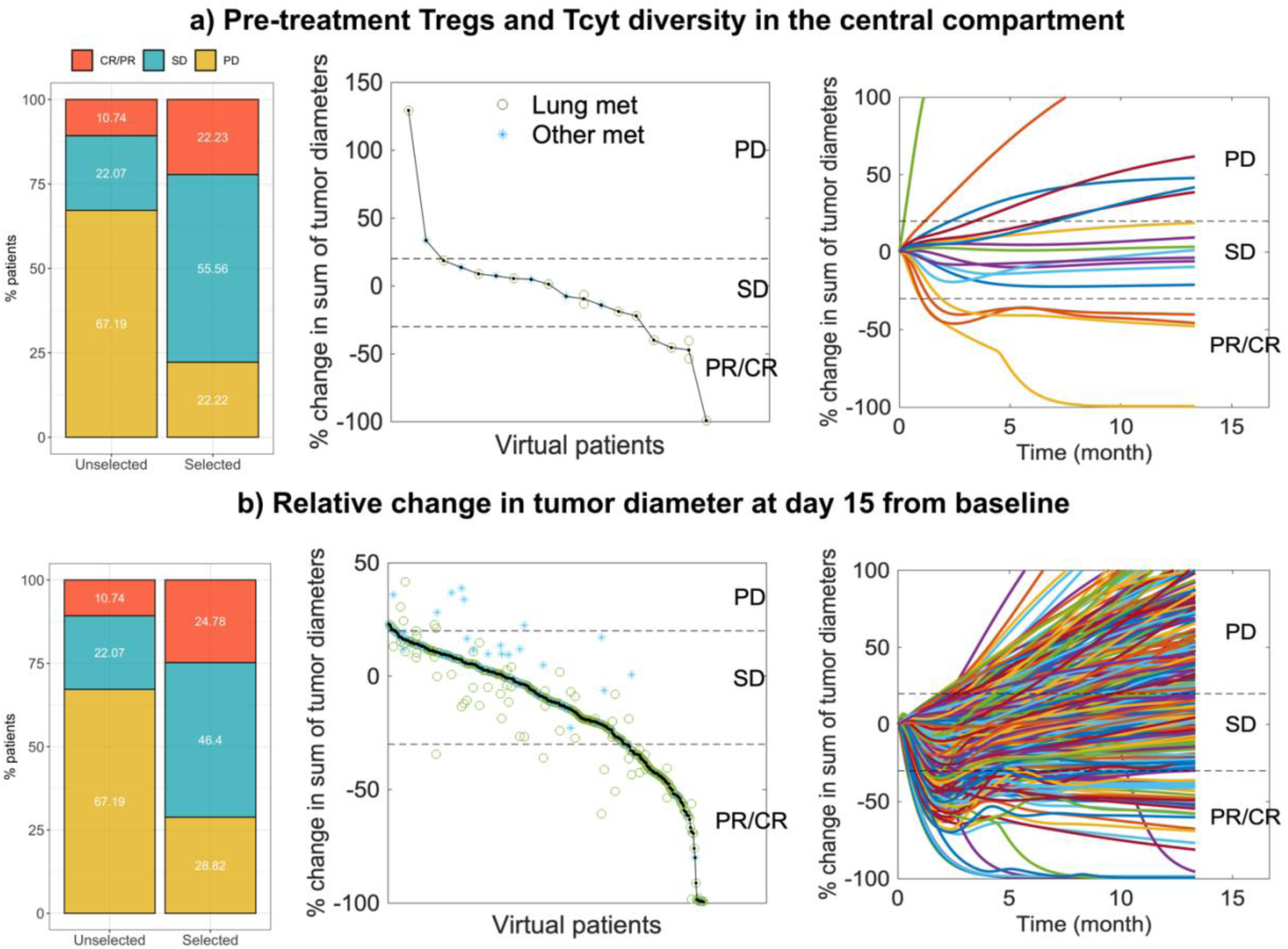
In-silico trial for biomarker validation. **a)** Outcome of an in-silico trial with patients selected based on non-invasive and pre-treatment biomarker combination of density of Tregs and Tcyt diversity in the central compartment. **b)** Outcome of in-silico trials with patients selected based on relative change in tumor diameter at day 15 from baseline. From left to right, fraction of patients with different response statuses compared to in-silico trial with unselected virtual patient population, variant of waterfall plot and spider plot are shown. In the waterfall variant plot, black dots represent the percentage change in the sum of tumor diameters of individual patients. Green circles and blue stars represent the change in the diameter of individual lung and “other” metastatic tumors.

As a representative of high specificity and high sensitivity, we chose the single on-treatment biomarker, relative change in tumor diameter at day 15 from baseline which can be determined by radiologic measurements. Among 829 virtual patients in the unseen cohort, 347 patients (42%) satisfied this biomarker criteria (relative change in tumor diameter at day 15 < 0.056). In-silico trial of the selected patient cohort showed an ORR of 24.78%, a DCR of 71.18%, and a median duration of response 10.5 months **(Fig 8b)**. Among patients with CR/PR, 78% had a response duration >= 6 months. Traditional waterfall plots are shown in **Fig S12**. Collectively, the exploratory in-silico analysis suggests that non-invasive biomarker candidates such as a combination of Treg density and Tcyt diversity in blood and the relative change in tumor diameter from baseline have promising ability to select patients for PD-1 inhibition. This work has elucidated strategies that may improve the performance of biomarker candidates that prompt clinical validation.

## Discussion

Although response rate of patients with metastatic TNBC to single-agent ICIs is low, some patients do respond with durable benefit. Therefore, a biomarker strategy to prospectively identify patients with potential to respond to ICI monotherapy is an important strategy for treatment de-escalation. The modest performance of existing biomarkers and the lack of understanding of strategies to improve their performance remains a barrier to patient selection for immunotherapy^57^. Here, we leveraged partially synthetic data generated from a multi-scale mechanistic QSP model, for an in-depth analysis of predictive biomarkers and to elucidate ways to improve their ability to accurately select patients with metastatic TNBC for PD-1 inhibition. In-silico patient-derived data employed in this study includes population distribution as well as longitudinal patient-level data. The in-silico approach undertaken overcomes challenges due to the limited availability of real-world patient-level data and enabled testing various what-if scenarios as an exploratory, hypothesis-generating analysis prior to clinical testing. For performance evaluation of biomarker candidates, we used two metrics: response probability and RIS. The former metric allowed us to explore the ability of biomarkers to achieve high specificity, while the latter metric enabled the identification of biomarkers that optimize the tradeoff between sensitivity and specificity. The biomarker candidates considered in this study are heterogenous (such as cell counts, cytokine concentrations, checkpoint expression) and require multiple assays. Thus, the main focus of the study was to identify the minimal set of biomarkers required, which reduces the need to perform several assays for biomarker quantification.

Results suggest that single pre-treatment biomarkers have limited sensitivity and specificity in patient selection, but pre-treatment biomarker combinations, such as the fraction of M1 macrophages among total macrophages in tumor combined with Treg/Tcyt ratio in lymph nodes possess a very high specificity. While pre-treatment biomarkers had limited ability to simultaneously achieve high specificity and sensitivity even when combined, on-treatment biomarkers, such as relative change in tumor diameter at day 15 from baseline and mature APC density in the tumor at day 15, were predicted to attain a better trade-off between sensitivity and specificity. Predictive biomarkers with high RIS can simultaneously achieve high specificity and sensitivity and are expected to have broader applicability as well as improved translatability across many patients. Among on-treatment biomarkers, relative change in biomarker candidate levels were superior in classification accuracy compared to on-treatment single time point measurements. However, on-treatment predictive biomarkers require the treatment to be initiated and thus might induce undesirable adverse events without any apparent efficacy for a subset of patients. Thus, in many cases, pre-treatment predictive biomarkers might be preferable compared to on-treatment biomarkers. Our findings demonstrating high specificity of pre-treatment biomarkers are promising and suggest that combining pre-treatment biomarkers is sufficient to achieve high response rates and disease control rates in clinical trials.

Our analysis also revealed several unexpected findings with implications for biomarker selection. While many biomarker candidates showed an increase in response probability and RIS after treatment initiation compared to baseline, this was not universally true for all biomarker candidates. Well-known biomarkers, such as PD-L1 in tumor and fraction of TILs, had reduced response probability and thus reduced specificity when tested after treatment initiation. When combinations of biomarkers were compared with single biomarkers, the extent of improvement in performance varied for different combinations. Unexpectedly, combinations of the best on-treatment biomarkers showed only modest improvement over best single biomarkers. Thus, biomarker candidates and the time point of measurement should be chosen carefully considering these observations.

In clinical practice, tissue-based biomarkers are most commonly used, but there are numerous efforts in identifying blood-based biomarkers to eliminate the need for invasive tissue biopsies^17^. To corroborate these efforts, we compared the performance of tissue-based biomarkers with blood-based biomarkers. Interestingly, our results also suggest that multiple blood-based pre-treatment biomarkers could be combined to achieve very high specificity. Many on-treatment tumor-based biomarkers had high RIS compared to blood or lymph node-based biomarkers. Unlike the measurement of cellular and molecular species in metastatic tumor samples, the relative change in tumor diameter at early time points of treatment initiation can be measured by imaging. This radiologic biomarker had the highest RIS among tested on-treatment biomarker candidates. These results are very promising and suggest that a high RIS implying a good balance between sensitivity and specificity can be achieved using non-invasive biomarker measurements. There is a need to select potential biomarkers and measurement time points on a case-by-case basis, accounting for factors such as measurement feasibility and desirable sensitivity-specificity trade-off to ensure that the benefits outweigh risks.

It should also be noted that the blood-based biomarkers tested only included T cell counts and ratios of cell counts. Liquid biopsy-based biomarkers such as circulating tumor cells and circulating tumor DNA have potential for predicting outcomes and recurrence risk in early-stage breast cancer^58, 59^. These biomarkers could also predict immunotherapy efficacy in metastatic cancer^17^, though this requires further investigation. Furthermore, our analysis was limited to cellular and molecular species explicitly included in the QSP model and thus did not include cell types such as B cells, NK cells, cancer-associated fibroblasts that might have an important contribution in determining the response status of patients receiving immunotherapy. Future extensions of the QSP model with additional species will evaluate their performance as predictive biomarkers. Our QSP model considered multiple tumors per patient to mimic the metastatic setting and we tested biomarker candidate levels averaged from tumors or lymph nodes in a patient. Due to inter-individual heterogeneity, variability in the responses of lesions within the same patient is inevitable^60^ and future work in developing a strategy for lesion-level biomarker analysis/response evaluation would be beneficial^61^. Biomarker candidates analyzed in this study are primarily non-spatial. Recognizing the importance of intra-tumoral heterogeneity in determining treatment outcomes^62^, imaging-based spatial biomarkers such as specific cellular neighborhoods have been identified in TNBC using multi-scale computational algorithms^63^. Using spatial QSP models that combine QSP with agent-based models to account for spatial heterogeneity^64–68^, it would be possible to extensively analyze the performance of spatial patterns as predictive biomarkers in metastatic TNBC.

Gene-expression profiling-based biomarkers have been identified to predict risk of recurrence or relapse and biomarkers predictive of treatment response in various cancer types^69–71^. ML models, such as neural networks and support vector machines, tend to have high accuracy in identifying prognostic or predictive biomarkers from very large datasets, but their applicability in clinical use was traditionally limited due to the black box nature of these models. Development of interpretable ML models can overcome these limitations and foster widespread clinical applications^72^. In this study, we used a cutoff-based biomarker testing algorithm combined with ML-based feature selection using random forest classifier. While ML-based feature selection improves computational efficiency, the cutoff-based biomarker testing provides information on the cutoff of biomarkers that maximizes the predictive power. A similar approach called PanelomiX^73^ has been implemented for generating biomarker panels where extrema of receiver operating characteristic curve are chosen as cutoffs and a random forest algorithm was employed to pre-filter features as well as cutoffs. This algorithm was implemented to predict the outcome of patients with aneurysmal subarachnoid hemorrhage^73^. Cutoffs are a key determinant that could tune the trade-off between sensitivity and specificity. These results emphasize that in addition to choosing the right biomarkers, choosing the right cutoff is also critical. There are multiple ways to choose cutoffs^73^ and generate biomarker panels that requires systematic exploration by future studies. Future studies analyzing the extent to which optimal cutoffs are cancer type/treatment specific would also be beneficial. While this study focused on investigating predictive biomarkers for pembrolizumab in the second or later-line treatment of metastatic TNBC, it is unclear if the findings and predictive biomarkers identified are treatment and cancer type specific. Nevertheless, the workflow established in this study is widely applicable. Future studies exploring predictive biomarkers for chemo-immunotherapy^74^ which is approved in the first-line setting of metastatic TNBC would be useful to inform treatment decisions.

A wide range of in-vitro, in-vivo and clinical data were utilized in model building, calibration, and virtual patient generation. Despite the use of multi-modal data, patient-level clinical data from patients treated with pembrolizumab monotherapy is extremely limited to validate the patient-level predictions of the QSP model. Thus, the QSP model generated longitudinal patient-level data, including the independent dataset utilized in this study, are partially synthetic, and therefore, caution is warranted before clinical applications. With the availability of more clinical data, it would be possible to further refine virtual patients generated using plausible ranges of cellular densities and molecular species concentrations^75^. Retrospective analysis of available biomarker candidate measurements including tumor volume measurements from previous clinical trials could be used to validate the findings of this study when patient-level data becomes available. Collectively, this study provides a wealth of testable hypothesis on strategies to improve patient selection as a first step towards precision medicine.

## Author contributions

Conceptualization: TA, RV, CAS-M, ASP. Methodology: TA, HW, RV, ASP. Investigation: TA. Visualization: TA. Supervision: RV, CAS-M, ASP. Funding acquisition: ASP. Writing—original draft: TA. Writing—review & editing: All authors

## Abbreviations

TNBC: Triple-negative breast cancer
TMB: Tumor mutational burden
TILs: Tumor infiltrating lymphocytes
dMMR: mismatch repair deficient
MSI-H: Microsatellite instability-high
TME: Tumor microenvironment
AI: Artificial Intelligence
QSP: Quantitative Systems Pharmacology
ML: Machine learning
CR/PR: Complete/Partial Response
SD: Stable Disease
PD: Progressive Disease
APCs: Antigen Presenting Cells
Tregs: Regulatory T cells
Tcyt: Cytotoxic T cells
LN: Lymph nodes
mAPC: mature Antigen Presenting Cells
Mac: Macrophages
Th: Helper T cells
PD-L1: Programmed cell death ligand 1
MDSC: Myeloid Derived Suppressor Cells
NO: Nitric Oxide
comp.: compartment
rel. change: relative change
UMAP: Uniform Manifold Approximation and Projection
ArgI: Arginase I
CCL2: CC motif chemokine ligand 2
IL-12: Interleukin 12
Immune supp.: Immunosuppressive
Immune act.: Immune activating
Ag: Antigen;
RIS: Responder Inclusion Score.

## Supplementary figures

**Figure S1:**
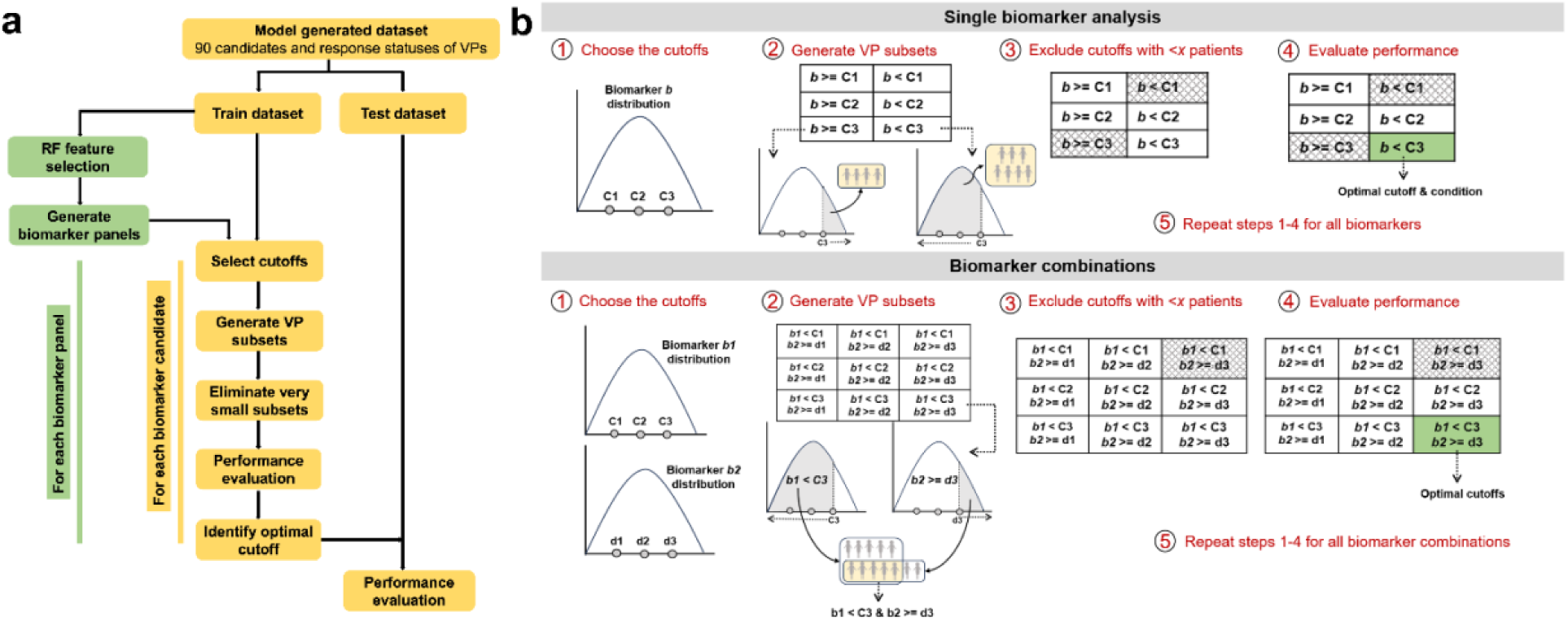
**a)** Outline of biomarker testing algorithm. Steps involved in single biomarker testing are shown in yellow. Modifications of the algorithm for testing biomarker combinations are shown in green. RF: random forest; VP: Virtual patients. **b)** Schematic illustration of the biomarker testing algorithm. *x* = 20 patients (∼2% of total) in this study.

**Figure S2:**
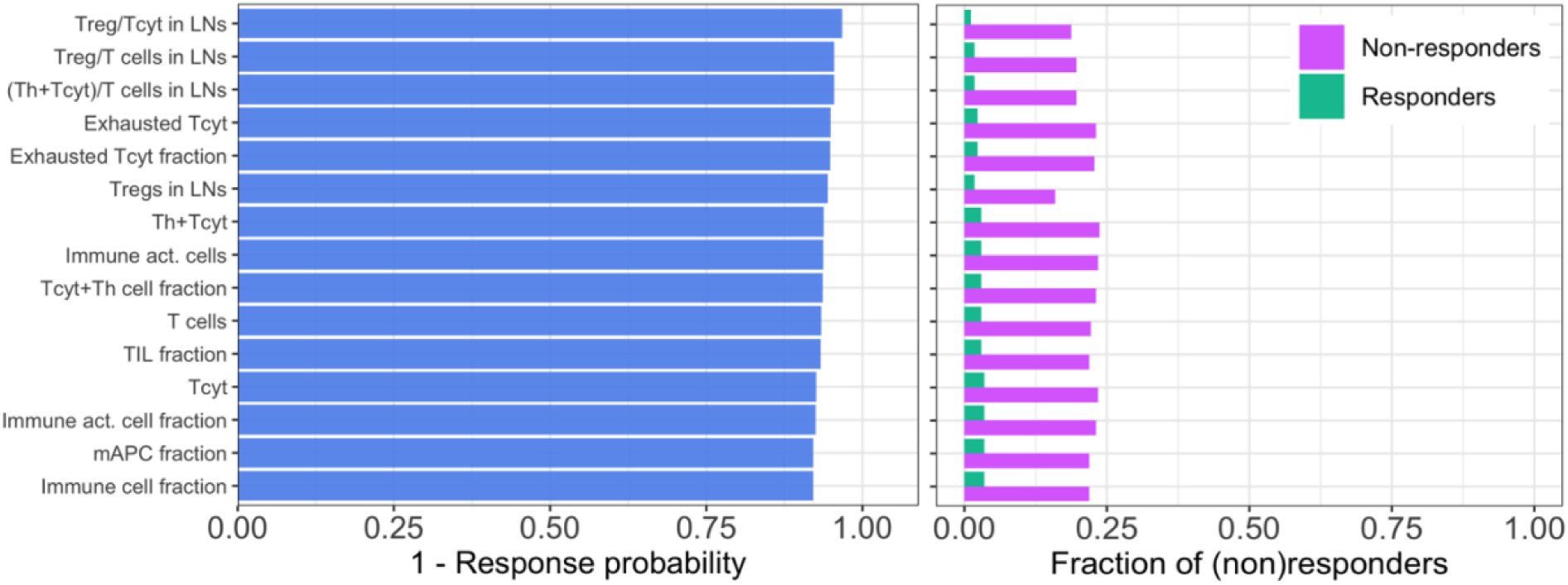
Performance measures of top 15 best single negative predictive biomarkers (pre-treatment). Treg: regulatory T cells; Tcyt: cytotoxic T cells; LN: lymph node; Th: helper T cells; act.: activating; TIL: tumor infiltrating lymphocytes; mAPC: mature antigen presenting cells.

**Figure S3:**
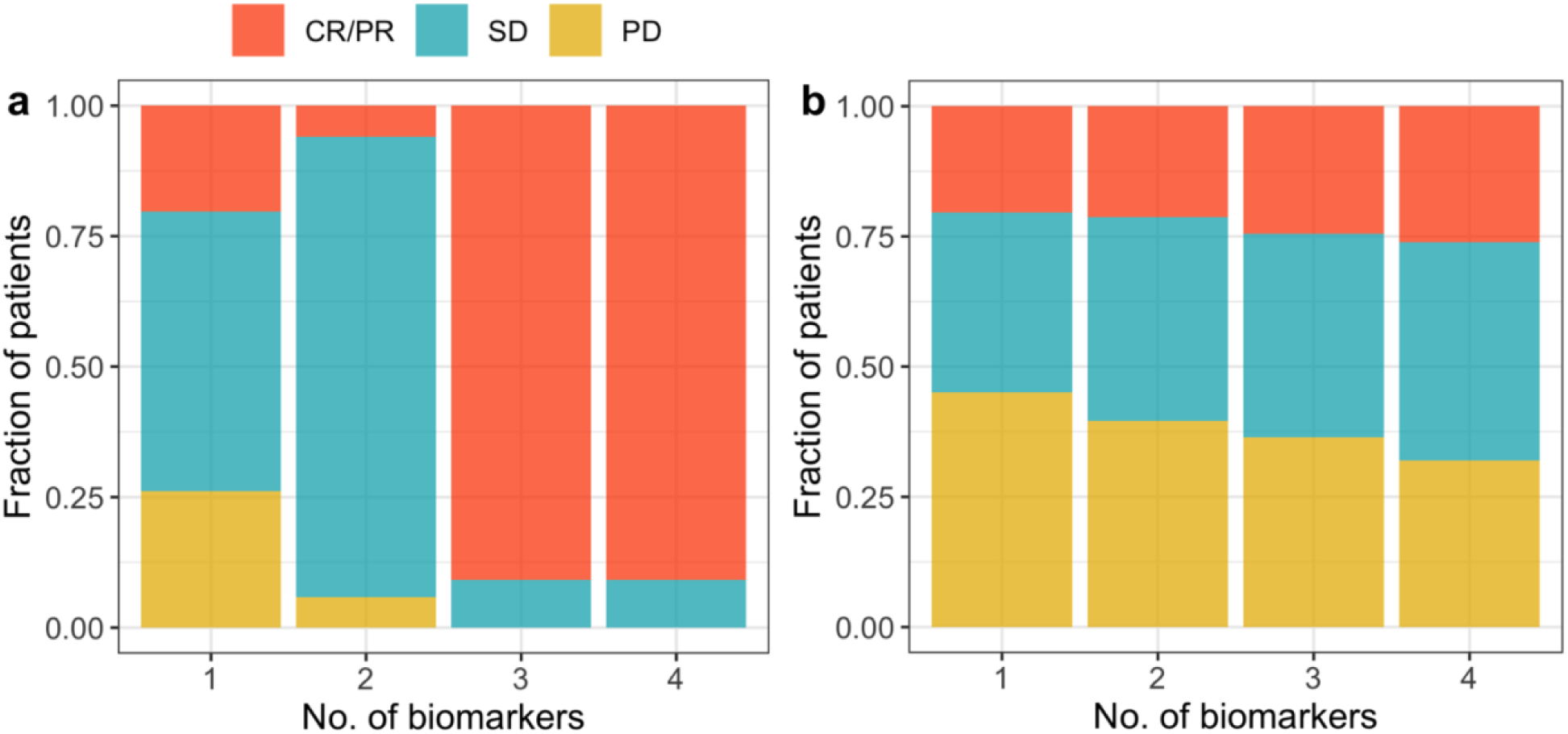
Fraction of patients with different response statuses within the subset of patients selected by best single and multivariate pre-treatment biomarkers based on performance measures: response probability **(a)** and RIS **(b)**. CR/PR: Complete/Partial response; SD: Stable disease; PD: Progressive disease.

**Figure S4:**
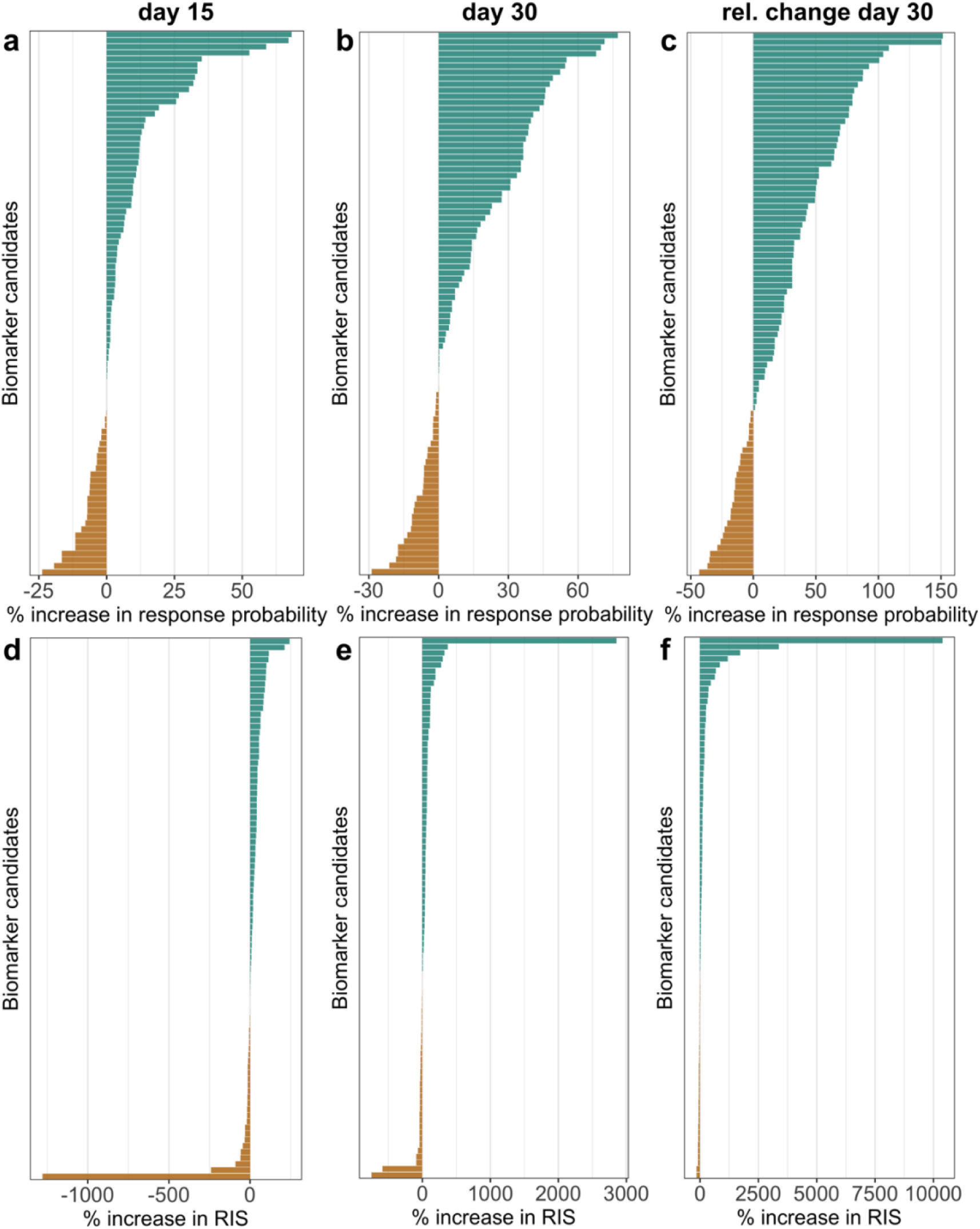
Percentage change in the performance of on-treatment biomarkers (day 15, day 30 and relative change on day 30 from baseline) with respect to performance of corresponding biomarkers at baseline. **a-c)** percentage change in response probability. **d to f)** percentage change in RIS. Biomarkers with increased and decreased performance after treatment initiation are depicted in green and golden yellow, respectively.

**Figure S5:**
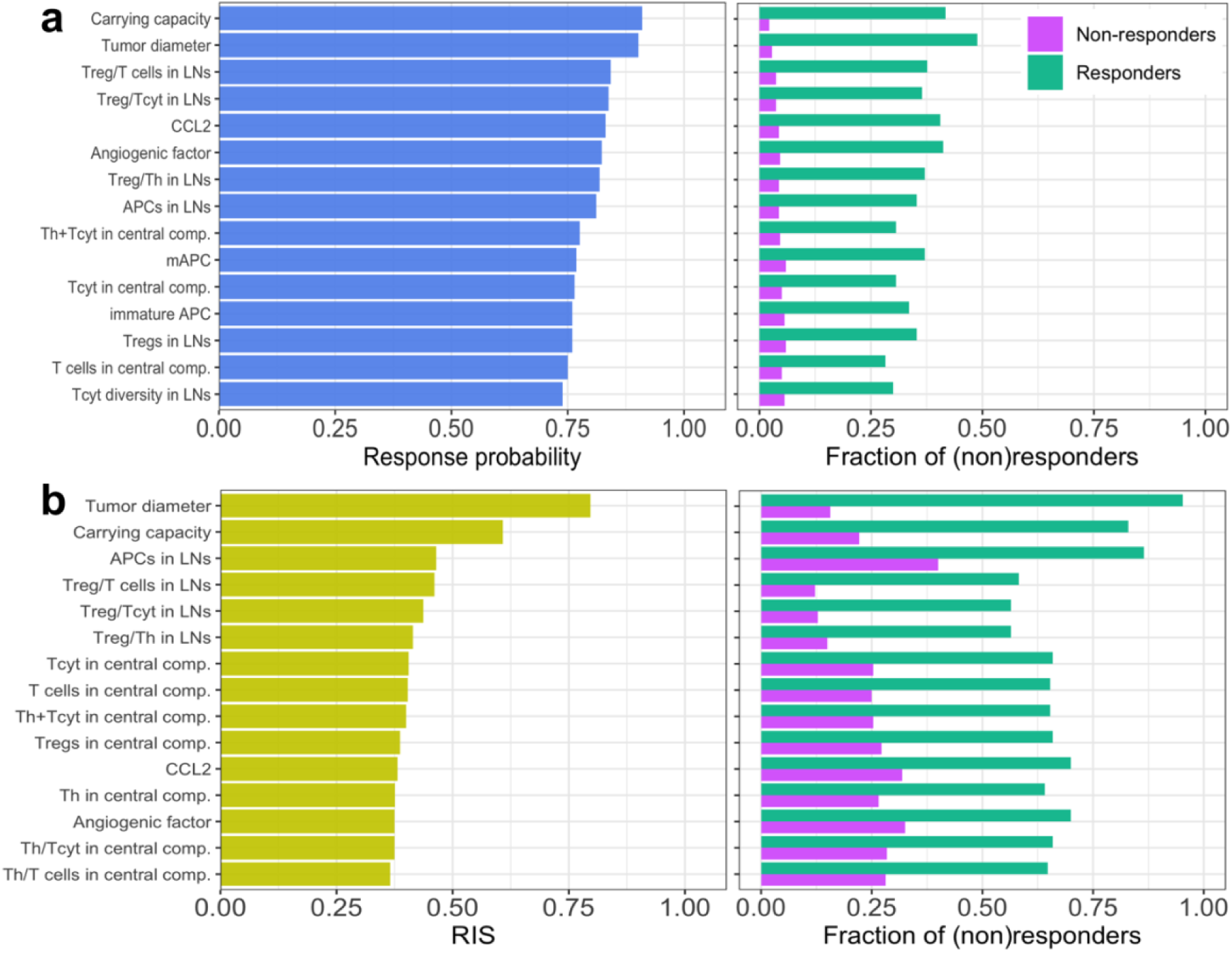
On-treatment biomarkers based on relative change on day 15 with respect to baseline. Top 15 best single biomarkers based on response probability **(a)** and RIS **(b)** are shown. Panels on the right show selected fraction of responders and non-responders from entire patient cohort by corresponding biomarkers. Treg: regulatory T cells; Tcyt: cytotoxic T cells; LN: Lymph node; Th: helper T cells; APC: antigen presenting cells; mAPC: mature antigen presenting cells.

**Figure S6:**
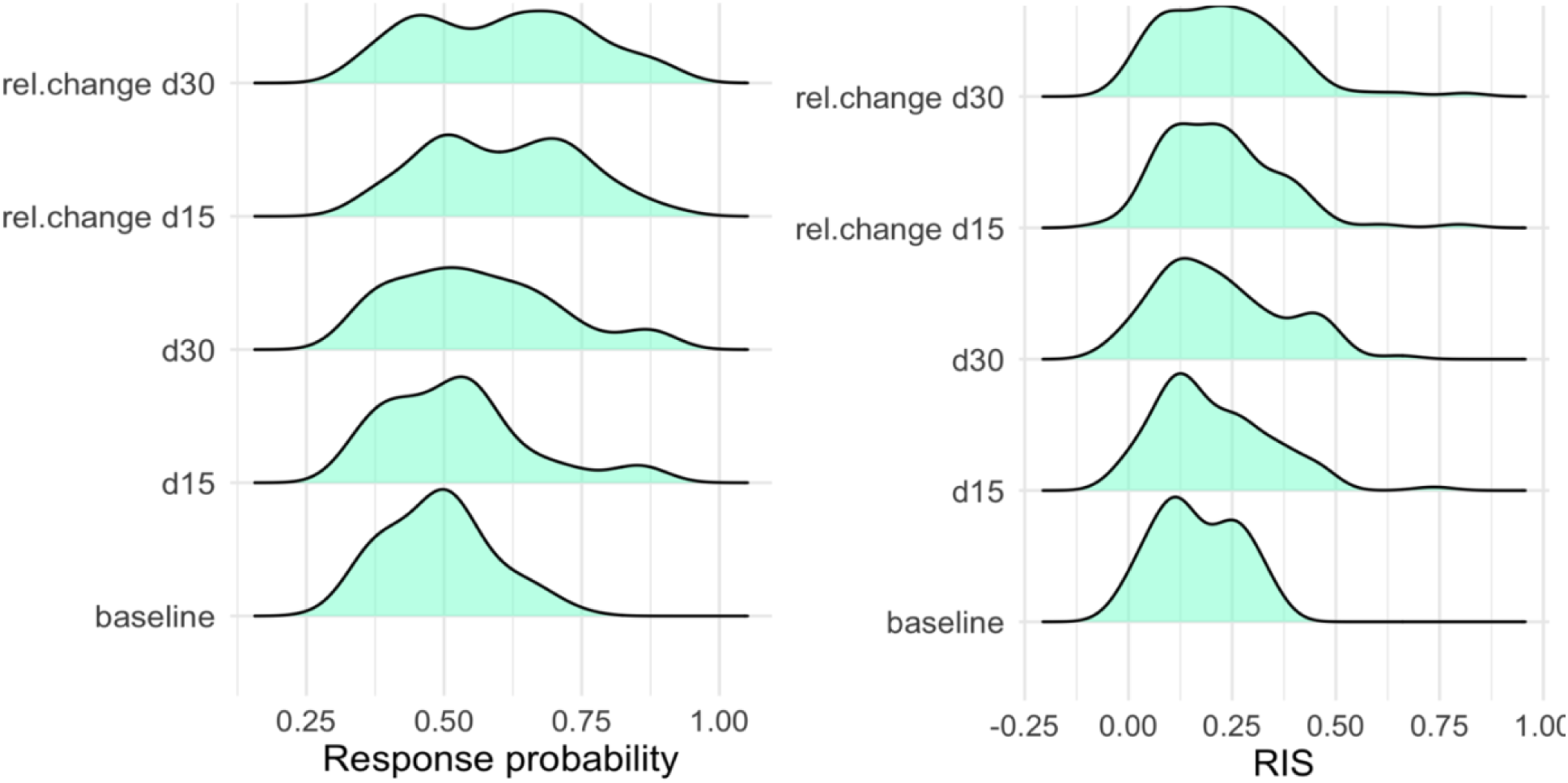
Distribution of response probability and RIS for pre-treatment and on-treatment single biomarkers. d15 and d30 represents biomarkers measured at single time points on day 15/30 after treatment initiation. “rel.change d15/d30” refers to relative change in biomarker levels on day 15/30 after treatment initiation with respect to the baseline.

**Figure S7:**
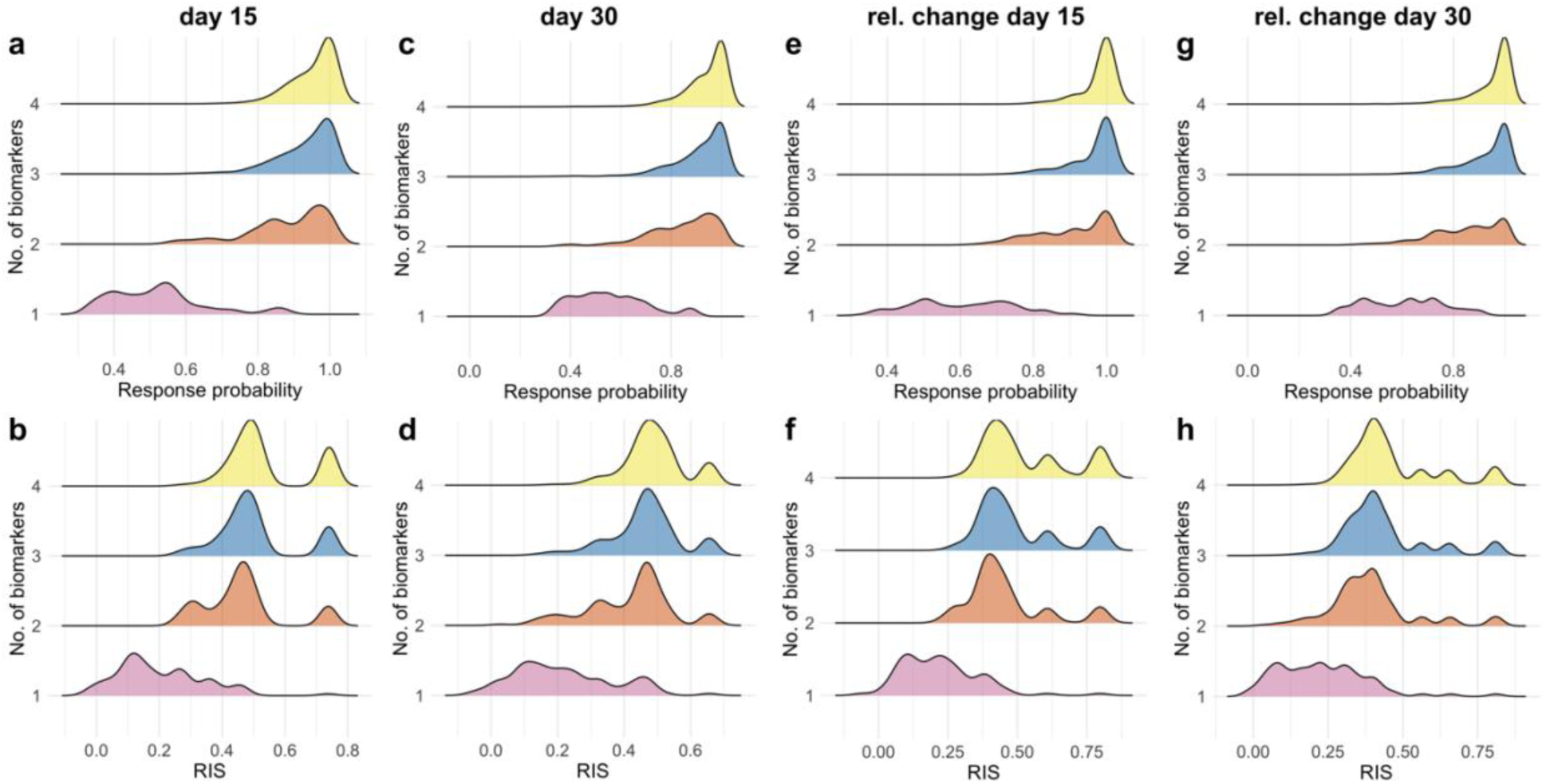
Distributions of response probability **(a, c, e, and g)** and RIS **(b, d, f, and h)** for single and multivariate on-treatment biomarkers. Colors distinguish single biomarkers from double, triple, and quadruple biomarker combinations.

**Figure S8:**
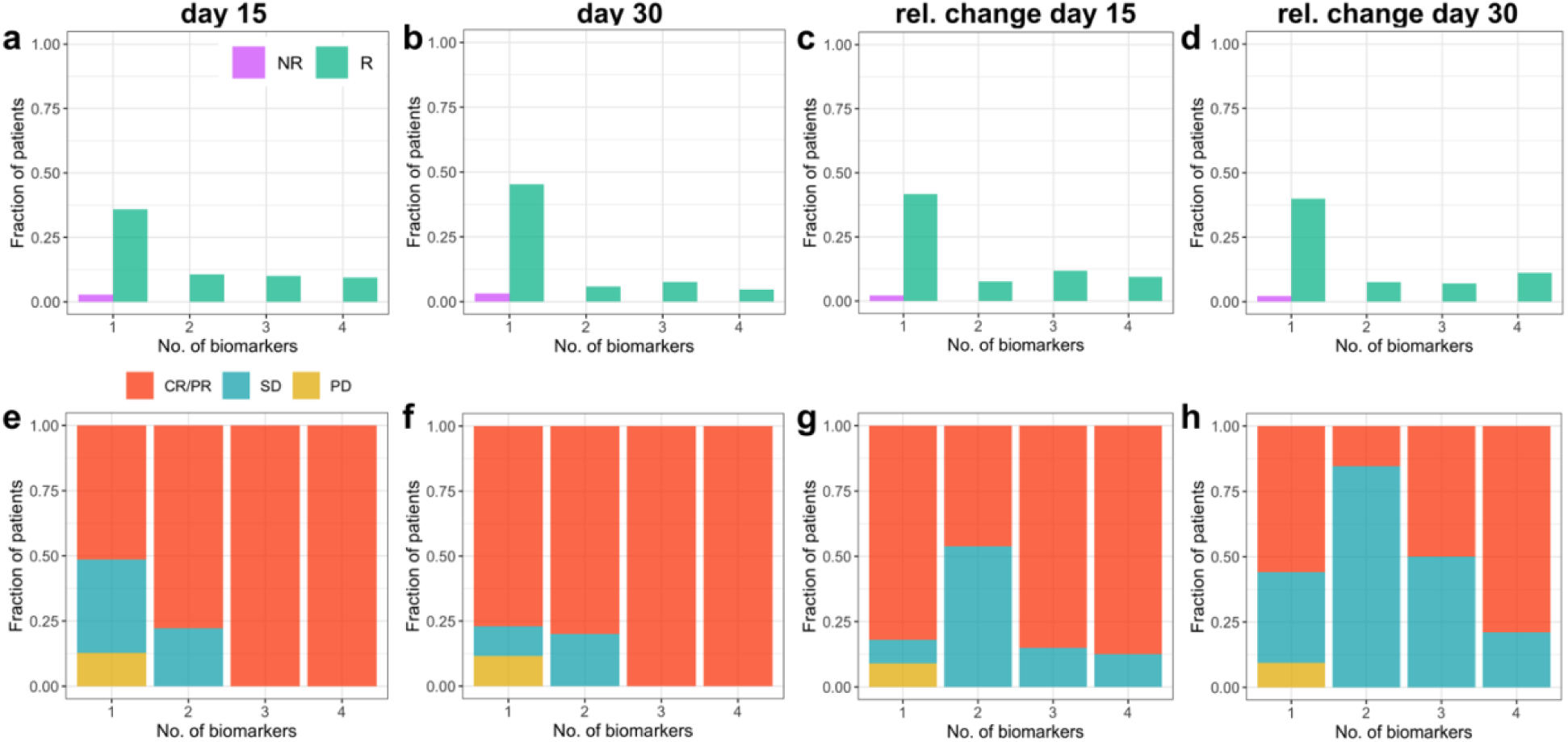
Composition of virtual patient subsets selected by the best single and multivariate on-treatment biomarker combinations based on response probability. **a-d)** Fraction of responders and non-responders from entire cohort selected based on biomarker levels at day 15 or day 30 and relative change on day 15 or day 30 from baseline. **e-h)** Fraction of patients with different response statuses within the subset of patients selected by the best single and multivariate biomarkers. CR/PR: Complete/Partial response; SD: Stable disease; PD: Progressive disease.

**Figure S9:**
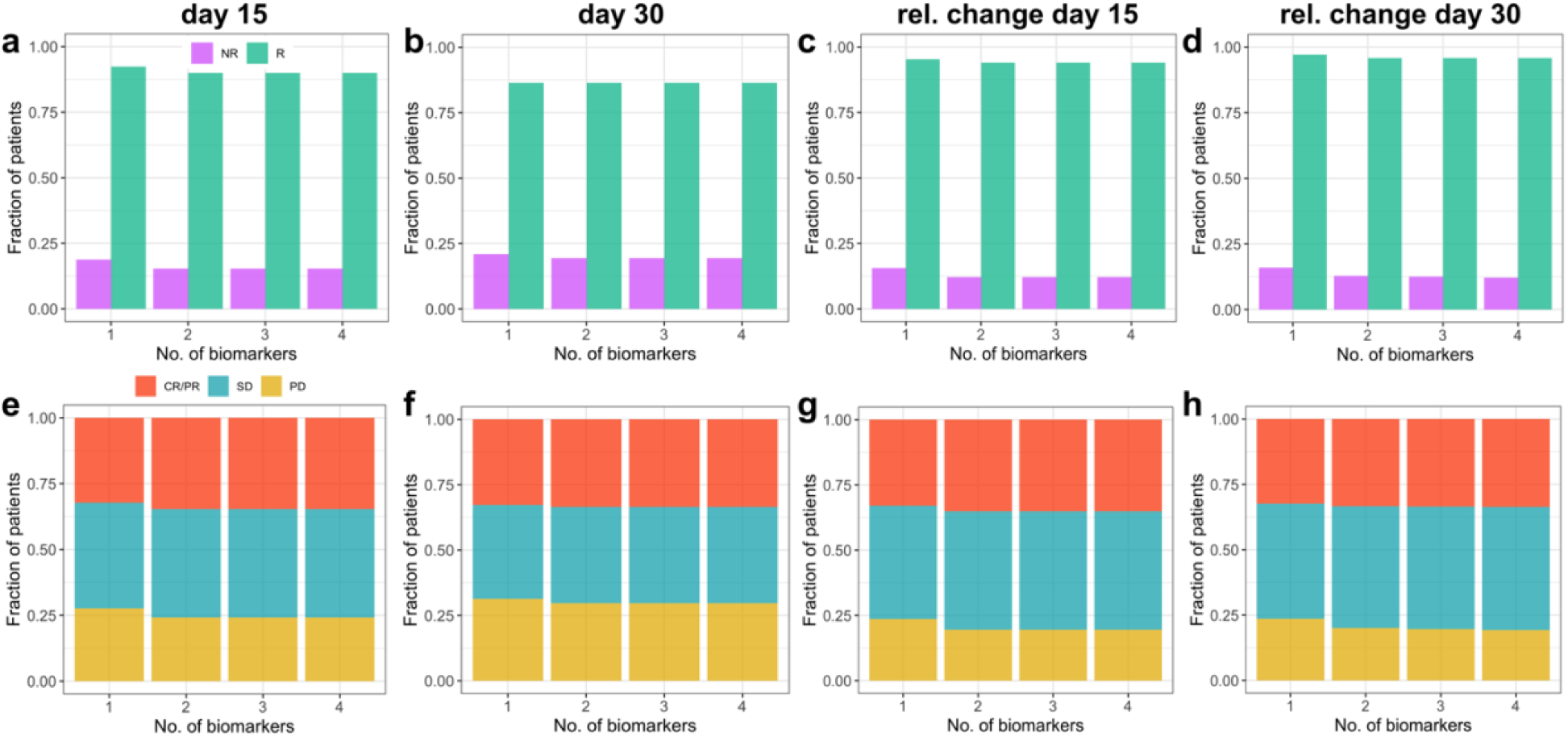
Composition of virtual patient subsets selected by the best single and multivariate on-treatment biomarker combinations based on RIS. **a-d)** Fraction of responders and non-responders from entire cohort selected based on biomarker levels at day 15 or day 30 and relative change on day 15 or day 30 from baseline. **e-h)** Fraction of patients with different response statuses within the subset of patients selected by the best single and multivariate biomarkers. CR/PR: Complete/Partial response; SD: Stable disease; PD: Progressive disease.

**Figure S10:**
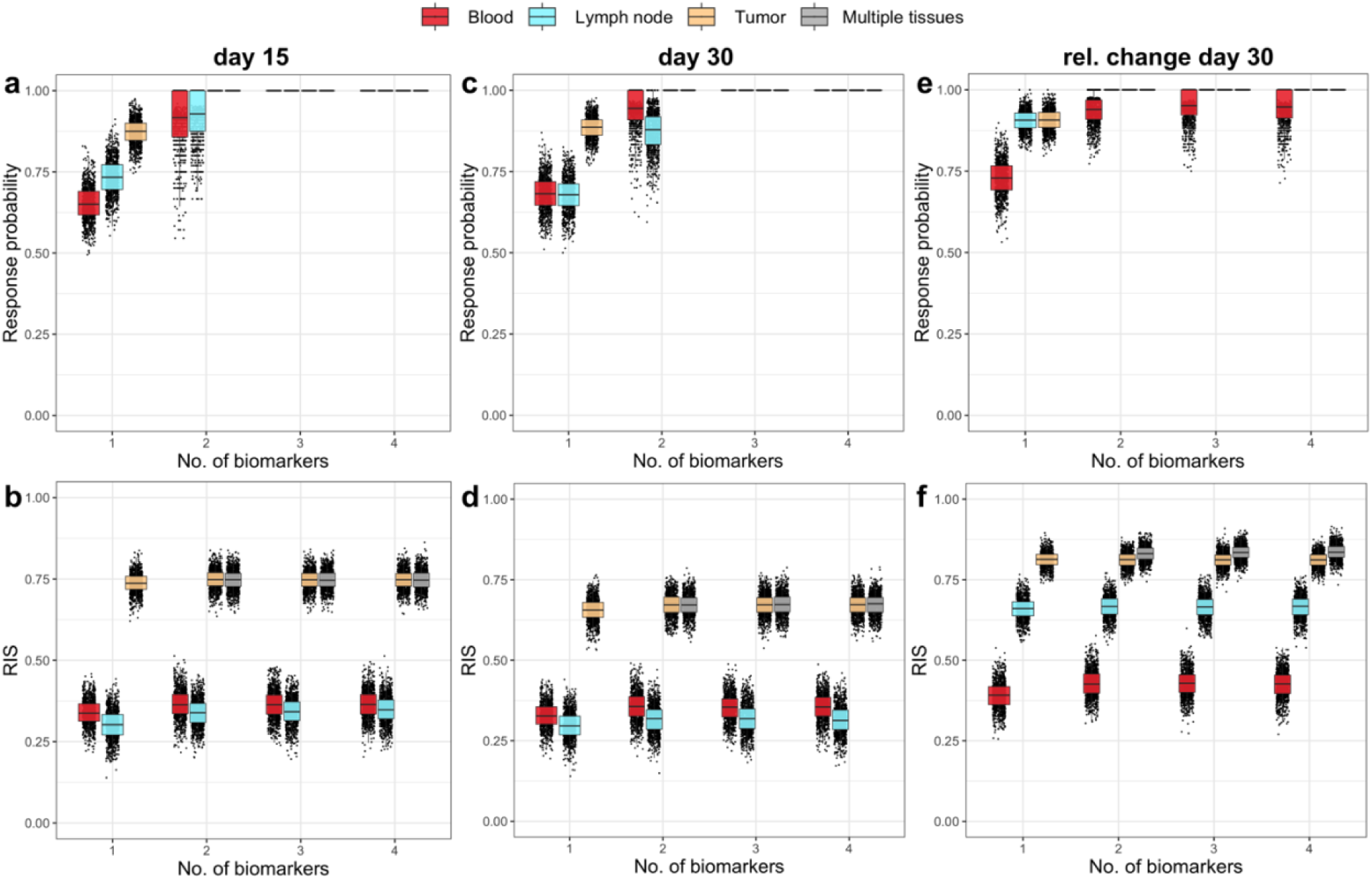
Response probabilities and RIS for on-treatment biomarkers from different compartments (blood, lymph node or metastatic tumor). **a and b)** single timepoint on-treatment biomarkers at day 15, **c and d)** biomarkers at day 30, **e and f)** relative change in biomarker levels at day 30 with respect to baseline. Colors indicate biomarkers/biomarker combinations from each compartment. Multiple tissues represent combinations of biomarkers from multiple compartments.

**Figure S11:**
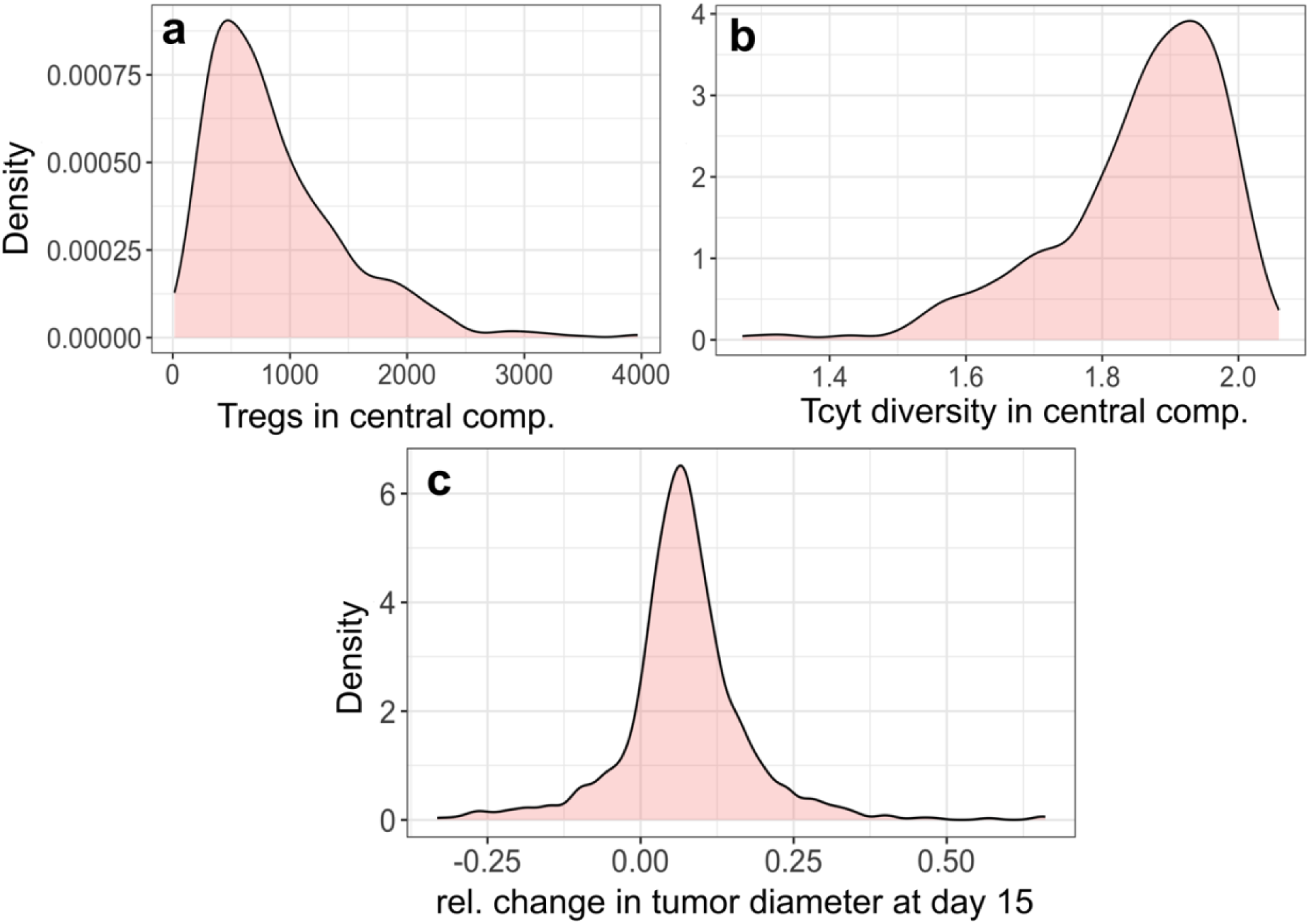
Distribution of biomarker levels in virtual patient cohort used for biomarker validation trials: **a)** pre-treatment density of Tregs in the central compartment in cells/ml, **b)** pre-treatment Tcyt diversity in the central compartment and **c)** relative change in tumor diameter at day 15 with respect to baseline (as fraction).

**Figure S12:**
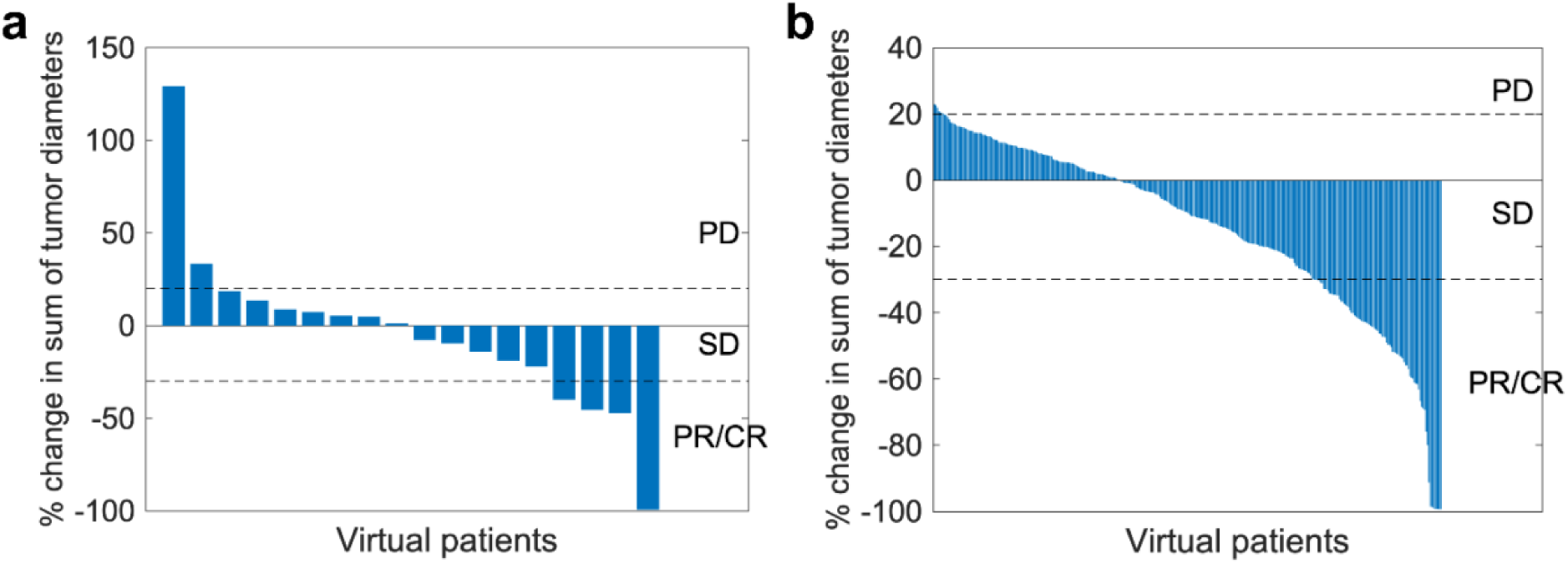
Traditional waterfall plots for in-silico biomarker validation trials shown in Figure 8. **a)** for in-silico trial with patients selected based on the density of Tregs and Tcyt diversity in the central compartment, and **b)** for patients selected based on the relative change in tumor diameter at day 15 from baseline.

